# The Inherent Flexibility of Receptor Binding Domains in SARS-CoV-2 Spike Protein

**DOI:** 10.1101/2021.08.06.455384

**Authors:** Hisham M. Dokainish, Suyong Re, Takaharu Mori, Chigusa Kobayashi, Jaewoon Jung, Yuji Sugita

## Abstract

Spike (S) protein is the primary antigenic target for neutralization and vaccine development for the severe acute respiratory syndrome coronavirus 2 (SARS-CoV-2). It decorates the virus surface and undergoes large conformational changes of its receptor binding domain (RBD) to enter the host cell, as the abundant structural studies suggest. Here, we observe Down, one-Up, one-Open, and two-Up-like structures in enhanced molecular dynamics simulations without pre-defined reaction coordinates. The RBD_A_ transition from Down to one-Up is supported by transient salt-bridges between RBD_A_ and RBD_C_ and by the glycan at N343_B_. Reduced interactions between RBD_A_ and RBD_B_ induce the RBD_B_ motions toward two-Up. Glycan shielding for neutralizing antibodies is the weakest in one-Open. Cryptic pockets are revealed at the RBD interfaces in intermediate structures between Down and one-Up. The inherent flexibility in S-protein is, thus, essential for the structure transition and shall be considered for antiviral drug rational design or vaccine development.

## Introduction

The severe acute respiratory syndrome coronavirus 2 (SARS-CoV-2) has caused over a 190 million infections and 4 million deaths, as of July 2021 (https://coronavirus.jhu.edu/map.html). It represents an urgent need for an effective medical intervention strategy, to avoid further social and economic consequences^1^. Different types of vaccines, for example, those from Pfizer-BioNTech or Moderna using the mRNA of Spike (S) protein, are currently available, and there are several FDA approved drug candidates under consideration^2,3^. At the same time, more infectious mutants such as B.1.617.2 (Delta) and B.1.427 / B.1.429 (Epsilon) have appeared, and some evade from the immune system^4,5,6,7^. Furthermore, the virus ability to infect a wide range of vertebrates, which could act as a reservoir, points out the future risk despite the vaccination progress^8^. A deeper understanding of the virus molecular structure and infection mechanism is crucial to stop the virus transmission including mutant strains^9^.

SARS-CoV-2, an enveloped positive single stranded RNA (+ssRNA) virus, has a large genome of approximately 30 kb encoding 29 proteins^9,10^. A transmembrane homotrimeric class I fusion glycoprotein decorating the virus, known as S-protein, plays a critical role in the viral cell entry^11,12^. In an immediate response to the pandemic, over a hundred Cryo-electron microscopy (EM) and X-ray crystal structures of S-protein have been reported and rapidly advanced in our understanding of the S-protein/receptor binding mechanism^13,14,15,16,17,18,19^. The N-terminal subunit (S1), which is composed of the N-terminal domain (NTD), the receptor binding domain (RBD) and two other subdomains (SD1 and SD2) (Figure S1)^16,20^, initially binds to the host cell receptor angiotensin-converting enzyme 2 (ACE2)^19^. This binding is followed by priming of the C-terminal subunit (S2) and its large conformational change leading to the membrane fusion for the cell entry^12,21^. S-protein is covered by 66 N-glycans, 22 per protomer, to evade from the host cell immune system^22,23^. To block the initial binding with ACE2 either by stimulating the immune system using vaccines or by neutralized antibodies or small-molecule drugs is the primary target for medical interventions. Numerous Cryo-EM structures reveal that the RBDs can take inactive Down and active Up forms including one-Up, two-Up and three-Up conformations (Fig. 1a)^13,14,15,16,17,21,24,25^. The RBDs are considered to undergo large conformational changes from the receptor inaccessible Down to the receptor accessible Up states in the ACE2 binding^16,19,26^. Note that most of the Cryo-EM structures representing the active Up forms were determined together with other proteins, such as antibodies or a fragment of ACE2 (Table S1). A recent single molecule fluorescence resonance energy transfer (smFRET) experiment suggested that the Down-to-Up transition occurs in S-protein even without its ligand, taking at least one transient intermediate^27^. This suggests the necessity to explore a wide conformational space of S-protein for describing transition pathways and intermediate structures. It would give us better understandings of the virus entry mechanisms and contribute to the development of antiviral drugs or antibodies.

**Figure 1.**
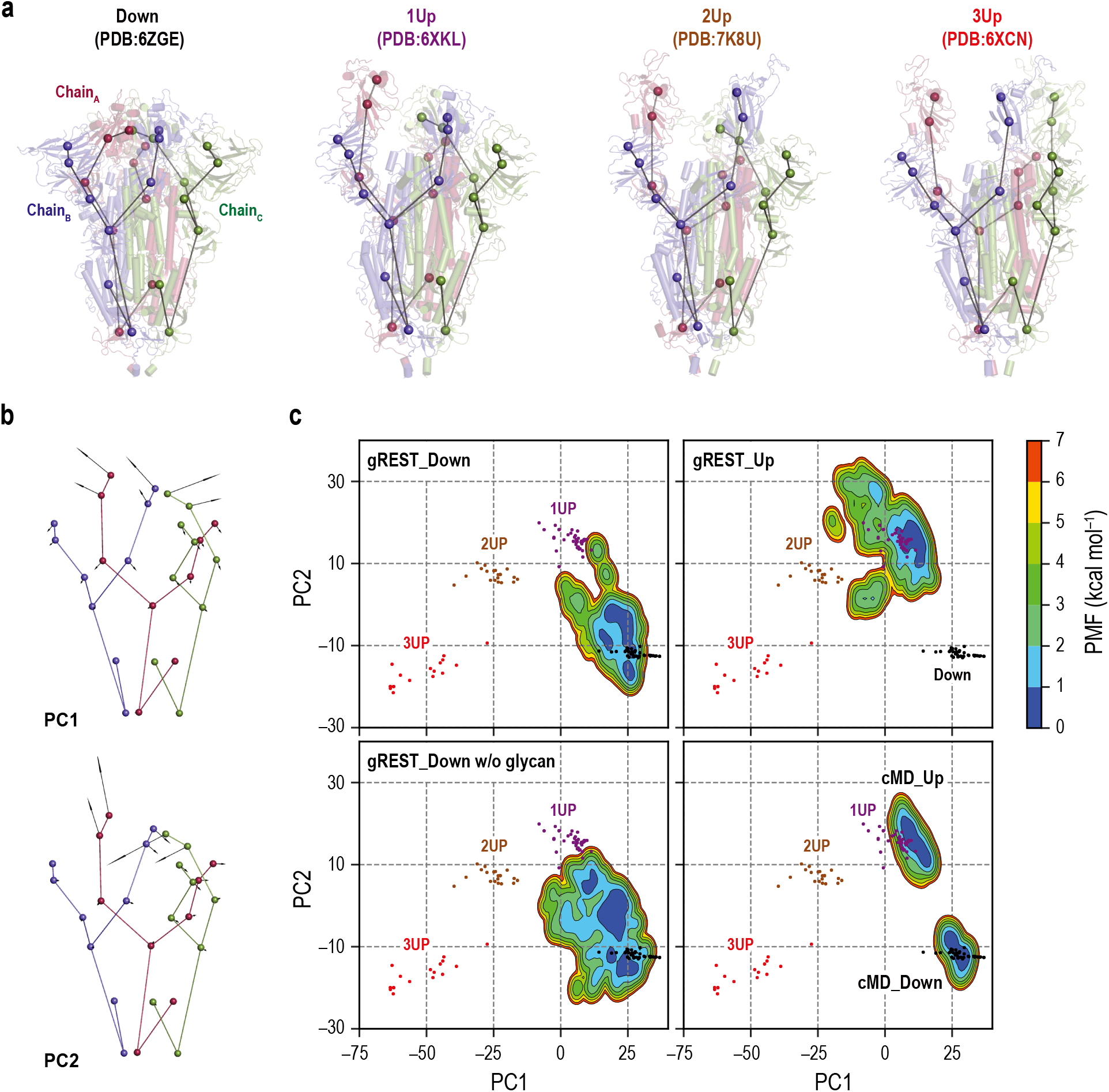
Comparisons of S-protein structures in MD simulations with Cryo-EM structures. **a)** 27 coarse-grained beads representations of four representative Cryo-EM structures: Down (PDB ID: 6ZGE), 1Up (6XKL), 2Up (7K8U), and 3Up (6XCN). Chain A, B, and C in S-protein are shown in red, blue, and green, respectively. **b)** The lowest two modes (PC1 and PC2) in principal component analysis of the 27 beads model of 126 Cryo-EM structures. PC1 and PC2 respectively represent a symmetric and an antisymmetric Down to Up motion of the receptor binding domains. The vectors are magnified 100 times for clarity. **c)** Free energy landscapes at 310 K along the PC1 and PC2 obtained from the simulations: (top) gREST_SSCR simulations with glycans starting from Down (500 ns) and Up (300 ns), (bottom left) gREST_SSCR simulation without glycans from Down (150 ns), (bottom right) conventional MD simulations with glycans starting from either Down and Up. The positions of Cryo-EM structures are also shown for comparison. Wherein, Down, 1Up, 2Up and 3Up conformations are shown in black, purple, brown and red dots, respectively.

Molecular dynamics (MD) simulations at the atomic level have been conducted to explore the conformational stability of S-protein by including the surface glycans, which are largely missed in the experimental structures. Several microsecond-scale MD simulations showed conformational flexibility of the stalk in S-protein, different levels of glycan-shielding between Down and Up, and their relevance in the ACE2 binding^22,28,29^. They also identified specific interactions between side-chain residues or those with glycans to stabilize either Down or Up^30,31^. The timescale of the Down-to-Up transitions in S-protein is far slower than that attainable one in the current MD simulations, leaving the inherent flexibility and transition pathways largely unknown. To overcome such difficulties, several enhanced sampling methods have been applied to investigate the Down-to-1Up-transition, including targeted MD^31^, steering MD^32^, nudged elastic band/umbrella sampling^33^, adaptive sampling,^34^ and weighted ensemble methods^35^. In most of these simulations, pre-defined reaction coordinates and/or bias potentials along the coordinates are used to enhance the motions of a single RBD in S-protein. Here we apply the generalized replica exchange with solute tempering of selected surface charged residues (gREST_SSCR)^36^, which enhances domain motions of a protein by exchanging the solute temperature of selected surface residues. gREST_SSCR is distinct from the other simulations applied to S-protein in that no reaction coordinates as well as bias potentials are used, allowing us to examine the inherent flexibility of S-protein involving more than one RBD. As listed in Table 1, the RBD motion in the RBD/SD1 monomer, the trimeric S-protein in the presence and absence of glycans, is systematically investigated. The intrinsic flexibility of S-protein observed in the enhanced sampling MD simulations suggests new mechanisms underlying the Down-to-Up transitions, glycan shielding for the binding with antibodies, and unprecedented cryptic pockets in intermediate structures during the transitions.

**Table 1.**
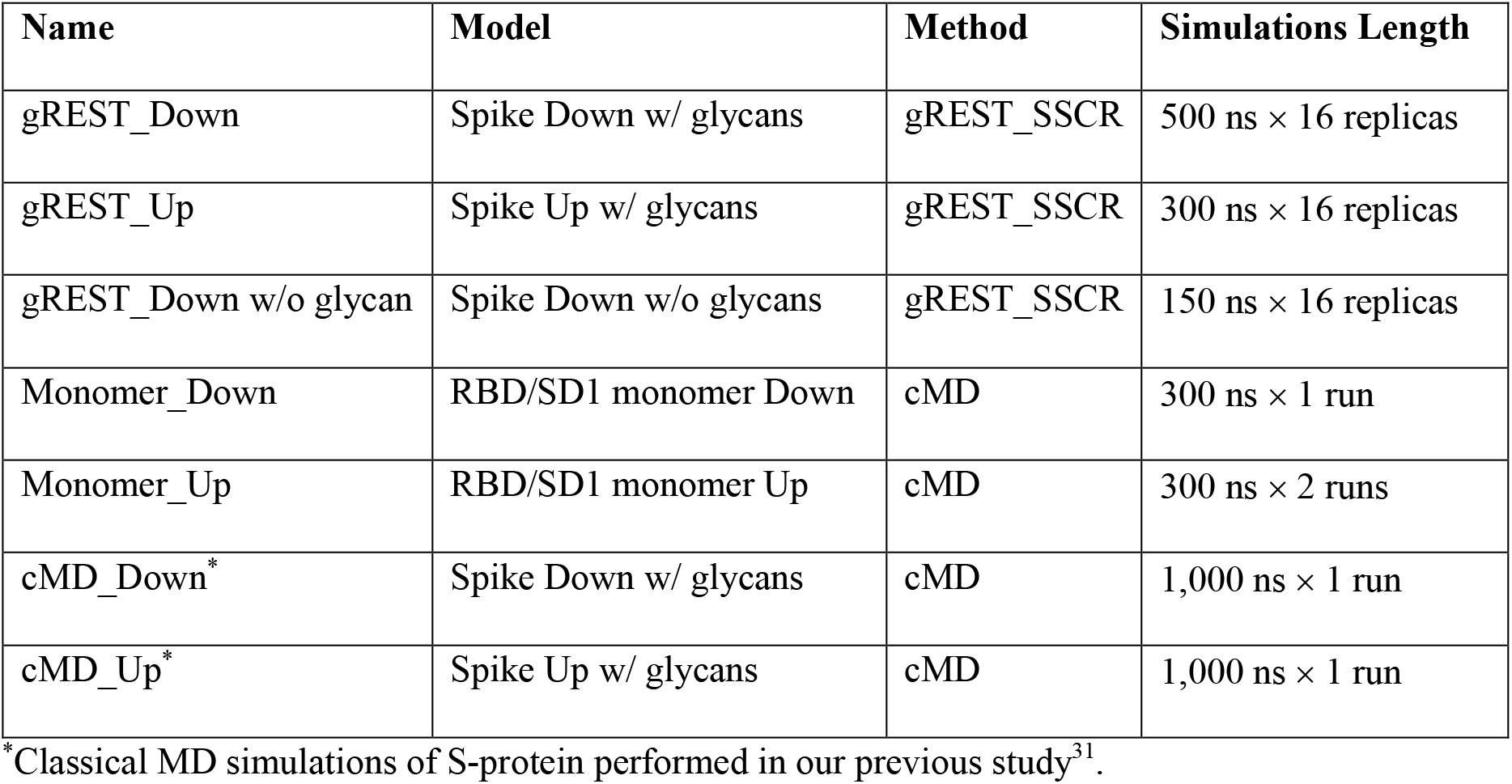
MD simulations of S-protein performed in this study.

| Name | Model | Method | Simulations Length |
| --- | --- | --- | --- |
| gREST_Down | Spike Down w/ glycans | gREST_SSCR | 500 ns × 16 replicas |
| gREST_Up | Spike Up w/ glycans | gREST_SSCR | 300 ns × 16 replicas |
| gREST_Down w/o glycan | Spike Down w/o glycans | gREST_SSCR | 150 ns × 16 replicas |
| Monomer_Down | RBD/SD1 monomer Down | cMD | 300 ns × 1 run |
| Monomer_Up | RBD/SD1 monomer Up | cMD | 300 ns × 2 runs |
| cMD_Down* | Spike Down w/ glycans | cMD | 1,000 ns × 1 run |
| cMD_Up* | Spike Up w/ glycans | cMD | 1,000 ns × 1 run |

\*Classical MD simulations of S-protein performed in our previous study<sup>31</sup>.

## Results

### Enhanced RBD motions in gREST_SSCR simulations

Table 1 lists all simulations performed in this study, wherein three gREST_SSCR simulations are involved. Two starting with a Down Cryo-EM structure (PDBID: 6VXX^20^) in the presence (gREST_Down) and absence (gREST_Down w/o glycan) of glycans and one from one-Up structure (PDBID: 6VYB^20^) (gREST_Up). All were solvated and ionized after modeling missing residues (SI Methods, Figures. S1 and S2). The simulations were performed using GENESIS software (2.0 beta)^37^, which was designed to achieve high performance and scalability on the supercomputer Fukagu^38^. In the solvated S-protein system including about 655,000 atoms, 2,048 nodes on Fugaku were used to achieve the performance of around 52 ns/day. The total simulation times including all the replicas are 8 *μs*, 4.8 *μs* and 2.4 *μs* in gREST_Down, gREST_Up and gREST_Down w/o glycan, respectively.

gREST_SSCR facilitates conformational sampling of a multi-domain protein by scaling the Coulomb and Lennard-Jones interactions of selected charged residues at domain interfaces through the solute temperature exchanges^36^. Here, 8 pairs of positively and negatively charged residues (870 atoms in total) in the RBD region were selected as the solute region of gREST_SSCR (Figure S3a). 16 copies of each simulation system were used to cover the solute temperature range from 310 to 545 K. For this method to work, the solute temperatures between neighboring replicas must be exchanged frequently. This, indeed, happened in all three simulations, showing sufficient overlaps of the potential energy distributions between neighboring replicas (Figure S3). All simulations show reasonable solute temperature exchange with the average acceptance ratio of 21.2%, 20.3% and 20.8% for gREST_Down, gREST_Up, and gREST_Down w/o glycan, respectively. The SD1/RBD hinge angle and the Cα root mean square deviation (RMSD) of each RBD upon fitting S2 provide simple measures of the Down-to-Up transitions: the RMSD and hinge angle in Up take the values of about 20 Å and 150°, respectively, while the hinge angle in Down is about 116°. Since gREST_SSCR is free from any pre-defined reaction coordinates, the Down-to-Up transitions happened not only in RBD_A_ but also in RBD_B_ or RBD_C_. Even two-Up-like conformations were observed in some replicas within 500 ns/replica (for instance, replica 8 in gREST_Down, replica 4 in gREST_Up, and replica 16 in gREST_Down w/o glycan) (Figure S4). Despite the large-scale motions, intra-domain structures of three RBDs and NTDs were kept stable. The Cα RMSD of RBD and NTD upon fitting to their own structures at 310 K in gREST_Down reveal about 1.4 and 2.2 Å, respectively (Figures S5a and S5b). They are comparable to those in the conventional MD^31^ (1.4 and 1.6 Å for RBD and NTD). Slightly larger RMSD values of NTD in gREST_SSCR are attributed to the loop regions abundant in NTD, as indicated in root mean square fluctuations (RMSF) (Figure S5c and S5d).

### Comparison between Cryo-EM and simulated structural ensembles

As of December 2020, 126 Cryo-EM structures of S-protein, including all Down, one-Up, two-Up and three-Up conformations, were deposited in the Protein Data Bank (Table S1). To describe the Down-to-Up transitions found in the Cryo-EM structures, we introduced three computational techniques: (i) the 9-beads representation per protomer as used in the studies by Henderson et al.^7,39^ to build up a 27-beads model of S-protein (Figure 1a and Table S2), (ii) the rotation scheme to make the most significant RBD motion always happen in ChainA (Figures S6 and S7), and (iii) the principal component analysis (PCA) on the 27-beads model of S-protein upon fitting all the beads to reduce the essential dimensions in the conformational space. In Figure 1b, the first principal component (PC1) represents a symmetric Down-to-Up motion involving all three RBDs, while the second component (PC2) reveals an asymmetric motion of RBDs where only RBD_A_ undergoes the Down-to-Up motion. The two lowest PCs cover about 85 % of the conformational fluctuations observed in the Cryo-EM structures. In Figure 1c, S-protein Cryo-EM structures with distinct RBD conformations, Down, one-Up, two-Up, and three-Up, are found in different clusters on the PC1-PC2 space (Figure 1c).

Next, we project the results of our previous 1*μs* conventional MD simulations (cMD_Down and cMD_Up) and the three gREST_SSCR simulations on the same space (Figure 1c). To obtain the distributions at 310 K in gREST_SSCR, we applied Multistate Bennett Acceptance Ratio (MBAR)^40^ for utilizing all the trajectories at different solute temperatures. The distributions of cMD_Down and cMD_Up overlap with the corresponding Cryo-EM structures (Down and one-Up, respectively), while there is a big gap between the two simulations. Instead, the distributions of gREST_SSCR simulations reach from the starting state to another: gREST_Down: from Down to one-Up, gREST_Up: from one-Up to two-Up, gREST_Down w/o glycan: from Down to one-Up. Due to the insufficient computational times, the initial structure dependences on distributions are found in all the three gREST_SSCR simulations. This issue would be resolved if we continue the simulations much longer timescales. More importantly, gREST_SSCR could cover more than one state, opening the possibility to investigate inherent flexibility of S-protein, transition pathways, and the intermediate structures. The distribution of gREST_Down w/o glycan is wider than that in gREST_Down, suggesting that glycans on the surface of S-protein play key roles in regulating RBD motions.

To focus on the Down-to-Up motions in S-protein, we define the hinge and twist angles using the Cα atoms in RBD and SD1 (Figure 2b). The hinge angle directly describes the Down-to-Up transitions, while the twist angle explains side motions of RBD. Larger values of the hinge and twist angles reveal the transition toward Up conformations. The gREST_SSCR distributions at 310 K on the hinge-twist angle space are compared to protomers of the 126 Cryo-EM structures in Figure S8, showing a good overlap between the Cryo-EM structures and structure ensembles at 310 K in gREST_SSCR as we see on the PC1-PC2 space. Interestingly, Monomer_Up, which consists of only a monomer containing SD1 and RBD (Up) sampled Down, one-Up, and one-Open states within 300 ns cMD simulations. The simulated ensembles in Monomer_Up is close to the Cryo-EM structures and the gREST simulations containing a whole S-protein.

**Figure 2.**
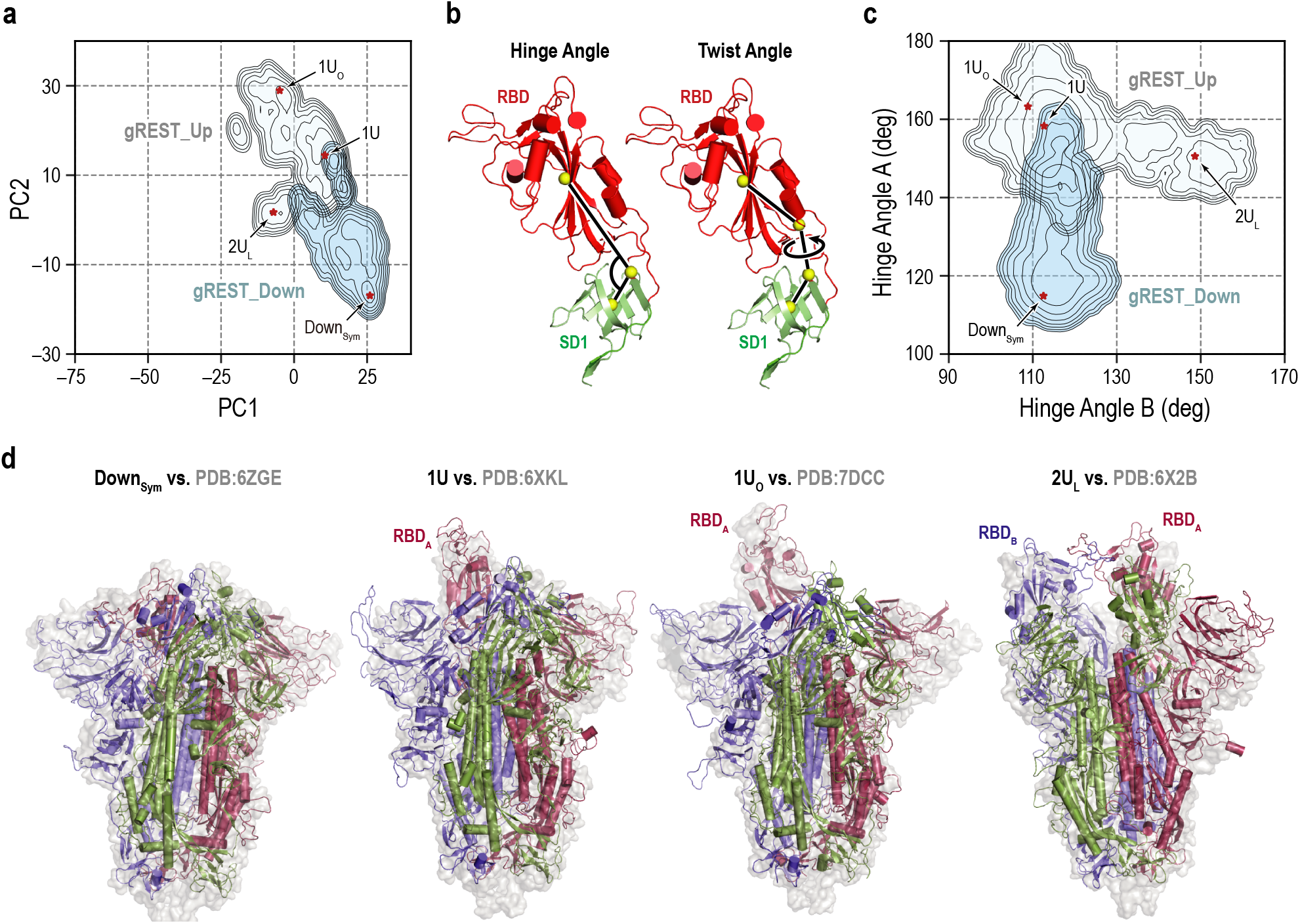
Representative RBD conformations from MD simulations. **a)** An overlay of the two free energy landscapes at 310 K along the PC1 and PC2 obtained from gREST_Down (light blue) and gREST_Up (light cyan) simulations. The red dots represent the positions of four representative RBD conformations: symmetric Down (Down_Sym_), 1RBD Up (1U and 1U_O_) and 2RBDs Up like (2U_L_) conformations. **b)** Definitions of the hinge and twist angles representing the RBD conformations. The hinge angle is determined by three center of masses (COMs, yellow spheres) of the core and top residues of SD1 (green, Cα atoms only) and the core residues of RBD (red, Cα atoms only). The twist angle is determined by the aforementioned COMs with an extra COMs of the bottom residues of RBD. **c)** An overlay of two free energy landscapes at 310 K along the hinge angles in RBD_A_ and RBD_B_ obtained from gREST_Down (light blue) and gREST_Up (light cyan) simulations. **d)** Four representative conformations from MD simulations (cartoon representation) in comparison with Cryo-EM structures (light grey surface). Chain A, B, and C in S-protein are shown in red, blue, and green, respectively.

On the Hinge_A_-Hinge_B_ space at 310 K, the Down-to-Up motion is also identified in ChainB in gREST_Up simulations starting from the one-Up structure in RBD_A_ (Figure 2c). We did not observe similar motions in the Hinge_A_-HingeC space, suggesting that the Up motions of RBD_B_ is followed from RBD_A_ to form two-Up structures. Taking together the information on the PC1-PC2 and the Hinge_A_-Hinge_B_ spaces, four major structures were identified: the Down symmetric (D_Sym_: (Hinge_A_, Hinge_B_) = (114.9°, 112.5°)), one-Up (1U: (158.3°, 112.7°)), one-Up-open (1U_O_: (163.3°, 108.9°)), and two-Up-like (2U_L_: (150.6°, 148.6°)). They are superimposed to the Cryo-EM structures having corresponding RBD conformations in Figure 2d. The simulation-derived D_Sym_ and 1U conformations are better aligned to the recent high-resolution structures of Down (PDB:6ZGE)^13^ and one-Up (PDB:6XKL)^14^, respectively. Intriguingly, RBD_A_ in 1U_O_ is well aligned to that in one of the RBDs in three-UP Cryo-EM structure (PDB:7DCC)^41^. RBD_A_ and RBD_B_ in 2U_L_ also agree well with those in two-Up Cryo-EM structure (PDB:6X2B)^39^, although 2U_L_ in gREST_SSCR remains inter-domain interactions between RBD_A_ and RBD_B_, which is completely lost in the two-Up Cryo-EM structures.

### The accessibility of RBD in different conformations

To get insights into the contribution of each conformation to the ACE2 and neutralizing antibodies (nAbs) bindings, the accessibility of RBD is examined in terms of the Solvent Accessible Surface Area (SASA)^30^. Figures 3a and 3b (and Figures S9-S11) show the per-residue SASA around the receptor binding motif (residues I410 to V510, referred to as RBM hereafter) calculated for D_Sym_, 1U, 1U_O_ and 2U_L_ conformations. Four mutational residues of concern, K417, L452, E484, and N501^5^, are highlighted in red. In Down conformation (D_Sym_), glycans at N165 and N343 largely shield RBM, leaving only small part accessible. RBM SASA increases in one-Up conformations (1U and 1U_O_) compared to Down, wherein 1U_O_ exhibits the utmost increase suggesting its potential contribution to the receptor and/or antibody binding. In contrast, the SASA increases only slightly, even decreases locally around residue C480, in two-Up (2U_L_) conformation due to the mutual interactions between two RBDs. This finding suggests that one-Up conformations are the primary target of nAbs and the bindings of multiple nAbs likely proceed in a stepwise manner through one-Up conformations. Notably, K417 and N501 are accessible only in one-Up conformations as expected, while E484 is accessible in both Down and Up, inferring a widespread effect of E484 mutations. On the other hand, L452 is not accessible in any conformations and hence is expected to rarely affect the binding with either receptor or antibody, in wild type.^42^ In contrast L452R mutation has shown to reduce neutralization^5^, suggesting its indirect contribution (e.g. the alternation of the S-protein conformational equilibria) to the immune escape. Figure 3c shows the calculated SASA for each epitope region of the four classes of nAbs (Class I: binds to Up for blocking ACE2, Class II: binds to Up and Down for blocking ACE2, Class III: binds outside RBM but recognize Up and Down, Class IV: binds Up without blocking ACE2)^5^. The epitope regions of Class I and II antibodies are largely exposed and almost completely de-shieled from glycans in Up conformations. On the other hand, the epitope region for Class III antibody is less exposed and largely shielded by glycans. This agrees with that Class III antibodies recognize “glycoepitopes”^43^. Note that the epitope region of Class IV antibody is slightly exposed only in 1U_O_ conformation. Further opening of RBD may allow the binding of Class IV antibody, such as CR3022, as suggested in a previous computational study^34^. Figure 3c shows the putative binding models with antibodies obtained by aligning the antibodies in Cryo-EM structures with each of Up conformations (see Figure S12 and S13 for detail). The 1U conformations can be targeted by any of Class I-III antibodies. On the other hand, 1U_O_ and 2U_L_ conformations are likely bound with Class I/II and Class II/III, respectively, suggesting the possible evade from some antibody attacks.

**Figure 3.**
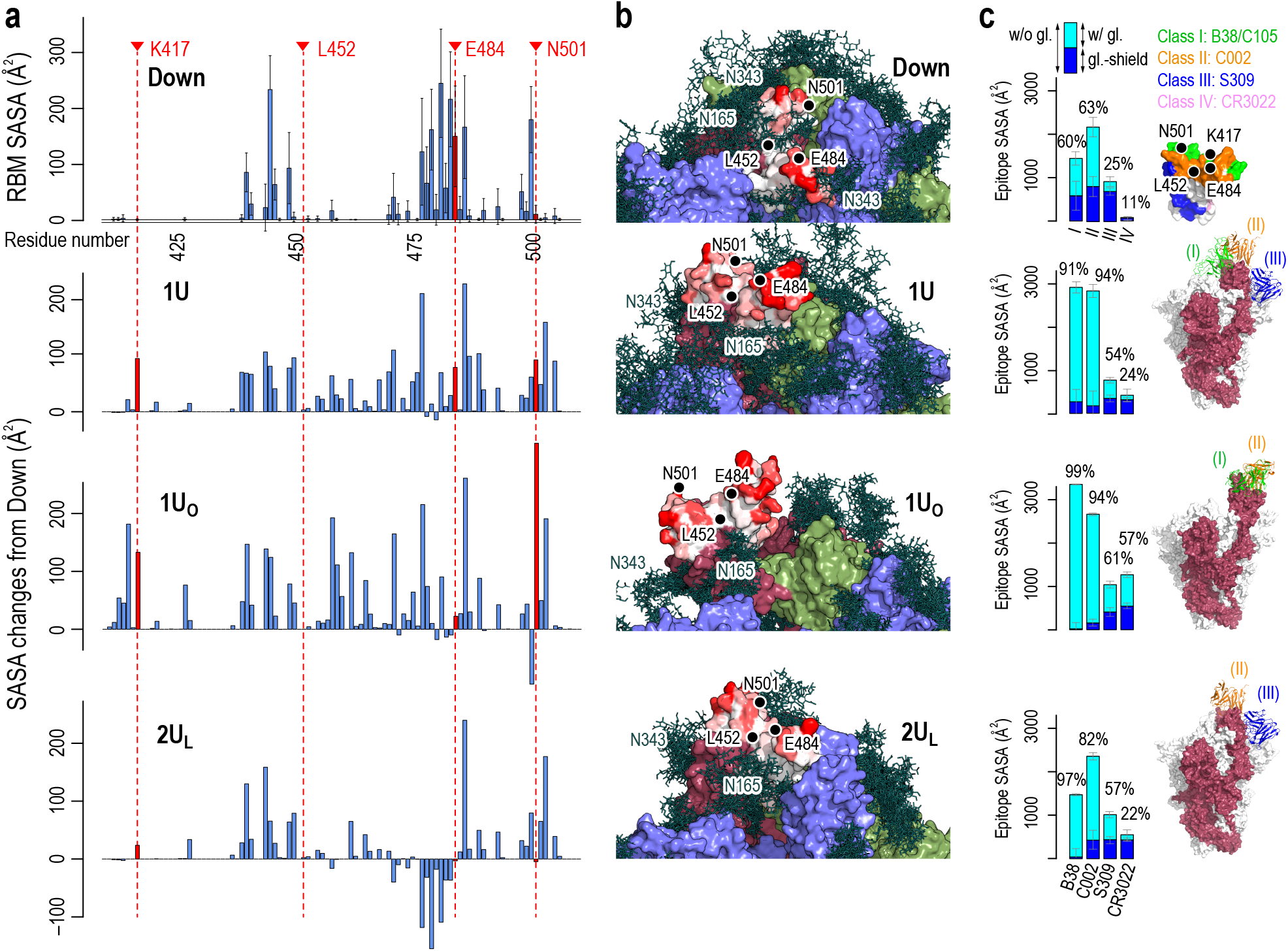
Accessibility of receptor binding motif (RBM). **a)** Per-residue solvent accessible surface area (SASA) values of the receptor binding motif (RBM, residues 410 to 510) in Down conformation (top) and their changes in Up conformations (bottom three). SASA values were calculated using the probe radius of 7.2 Å. Four mutational residues, K417, L452, E484, and N501, are highlighted in red. **b)** The surface representation of RBM SASA (white to red for 0 to 260 Å^2^, the maximum value in Down, the values higher than this are truncated for consistent color scheme). Chain A, B, and C in the protein are shown in red, blue, and green surfaces, respectively, while a collection of glycans from 10 snapshots are shown in stick representation. **c)** Epitope SASA and glycan shielding of four types of neutralized antibodies, B38 (Class I), C002 (Class II), S309 (Class III), and CR3022 (Class IV). Sum of SASA with glycans (cyan) and the glycan-shield (blue) gives SASA without glycans. The ratio of the SASA with glycan over that without glycans is shown. The right most column show the putative interaction models with three classes of antibodies: Class III (S309, PDBID: 6WPT, blue), Class II (C002, PDBID: 7K8T, orange), and Class I (C105, PDBID: 6XCM, green).

### Comparison between smFRET and simulated structural ensembles

Using smFRET, Lu et al. showed the conformational dynamics of S-protein in the presence or absence of its receptor, hACE2.^27^ They characterized four structural ensembles including two types of Down (major and minor), one intermediate and one-Up, suggesting the inherent flexibility of the RBD region regardless of the receptor binding. To examine whether gREST_SSCR could reproduce the experimental data, we first classified each gREST_SSCR trajectory at 310 K by means of *k*-means clustering and re-clustering guided by the hinge and twist angles distributions (Figures S14-S16, Table S3) and then computed the distance between residues 425-431 in RBD_A_ and residues 554-561 in SD1_A_ to correlate the smFRET intensity by Lu et al.^27^ By combining the distance distributions from gREST_Down and gREST_Up, we obtained the five conformational ensembles (Figure S17): Down symmetric (Down_Sym_), Down like (Down_Like_), two intermediates (Int2 and Int3), and Up (1Up). The smFRET distance alone cannot distinguish various Up conformations including 1U, 1U_O_, and 2U_L_ and thus we refer them to as 1Up. Down_Sym_ and Down_Like_ give distributions in the range of 30-35 Å, while 1Up shows around a median distance of 47 Å. The distributions of Int2 and Int3 are median distances of around 38 Å and 40 Å, respectively. Since Int3 has a large distance distribution that overlaps with Int2 and both intermediates might be indistinguishable in the smFRET experiment, the four conformations corresponding to the smFRET intensity of 0.8, 0.5, 0.3 and 0.1 in the absence of hACE2^27^ might be Down_Sym_, Down_Like_, Intermediate (Int2 and Int3) and 1Up. Each ensemble is also characterized with hinge angle distributions: Down_Sym_:Hinge < 120°, Down_Like_: Hinge < 130°, Int2:120° < Hinge < 140°, Int3:Hinge ~ 140°, 1Up:140° < Hinge < 160°.

### Transition pathways and transient interactions stabilizing the intermediate structures

Based on the free-energy landscapes at 310 K and the comparison between the gREST simulations and smFRET experiments, we now have evidence on the inherent flexibility of S-protein in the absence of its ligand. We next focus on molecular mechanisms underlying the Down-to-1Up transitions in S-protein in terms of the correlated motions of the hinge and twist angles in three RBDs (Figure S18). Note that such correlated motions are hardly obtained using targeted MD simulations (Figure S19) or the MD simulations enhanced with pre-defined reaction coordinates^35,44^. As expected, Hinge_A_ and TwistA are highly correlated with each other (Figure S18c), while Hinge_A_ and Hinge_B_ show almost no correlations (Figure S18a). Some correlations exist between HingeÅ and Hing_C_/Twist_B_, which appear to relate with the preference of two intermediates.

To examine the origin of such correlated motions in three RBDs, contact analysis (Figure S20) and hydrogen bond (HB) analysis (Figure S21) were carried out for each cluster observed in gREST_Down and gREST_Up simulations as we did in the previous study^31^. In Figure 4, several key contacts and hydrogen bonds are highlighted to see drastic changes of the interactions in the Down-to-1Up transition. The glycans at N343_B_ and N234_B_ switch the interactions with RBD_A_ along the transition from Down to one-Up-like. In the intermediates, I2a and I3a, the glycan at N343_B_ changes its contact partners by inserting underneath RBD_A_. Concurrently, the formation of salt-bridge interactions between RBD_A_ and RBD_C_, for instance, R408_A_-D405_C_ and R408_A_-D406C, lift up RBD_A_ from its Down to intermediate structures. Finally, in 1U_L_, N234_B_ inserts into the newly formed cavity between RBD_A_ and RBD_B_, forming new contacts with D428_A_ and T430_A_. The glycan at N165_B_ is in contact with RBD_A_ throughout the transition, likely acting as a main barrier for the transition.

**Figure 4.**
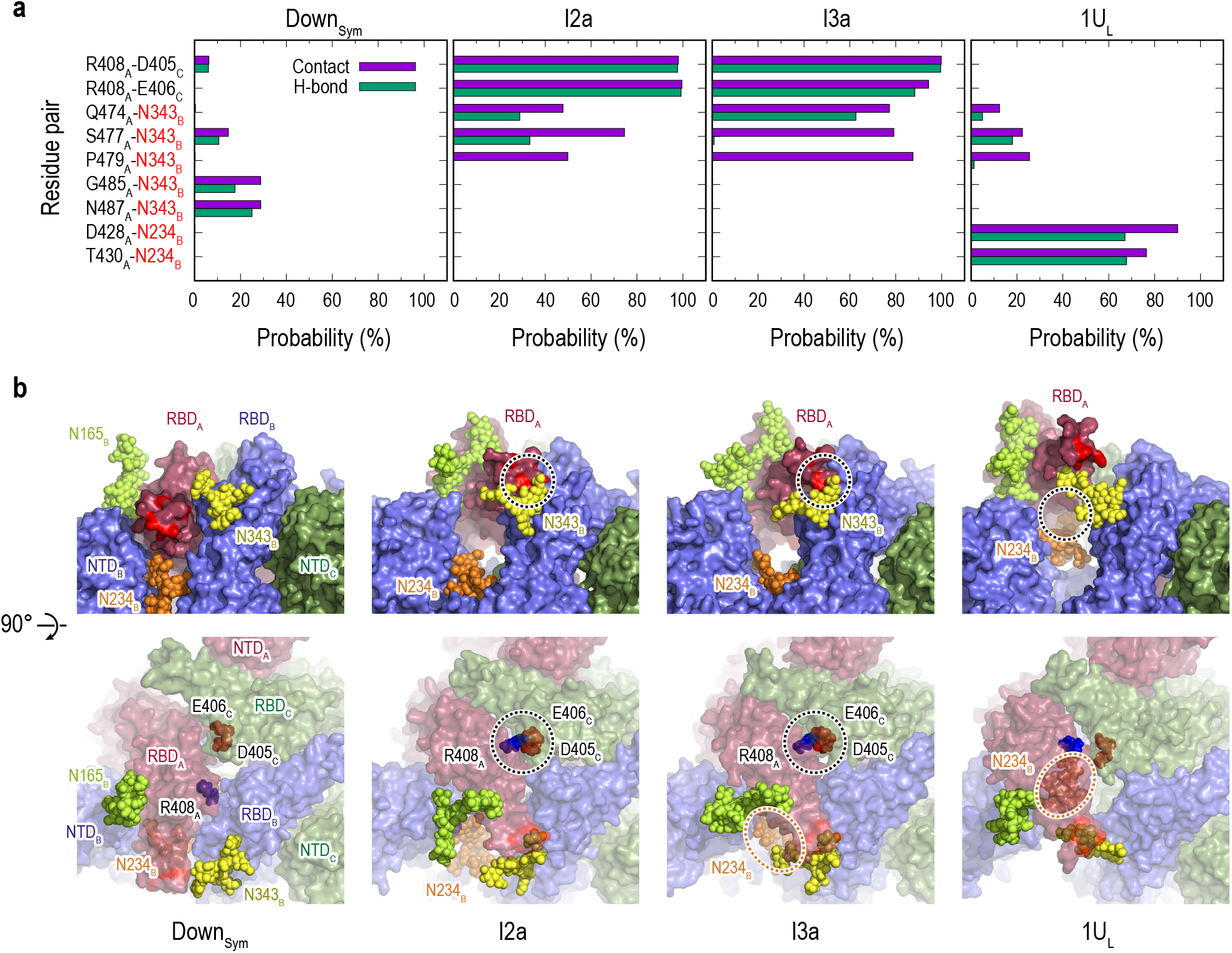
Protein-protein and protein-glycan interactions critical for Down-to-Up transition. **a)** Probability of finding the hydrogen bond (green) and contact (purple) pairs between protein residues or protein-glycans that markedly change along the transition pathway (D_Sym_, I2a, I3a and 1U_L_). All hydrogen bond (probability of finding of >50%) and contact pairs (probability of finding of >70%) are shown in figure S19 and S20, respectively. **b)** Typical snapshots of the protein-protein and protein-glycan interactions along the transition pathway. Chain A, B, and C in the protein are shown in red, blue, and green surfaces, respectively. Glycans at N165, N234 and N343 are shown with spheres in lime, orange and yellow color, respectively. The transient N343_B_–RBD_A_ contact are highlighted in red surface (top). The salt-bridges formed by R408_A_ (blue), E406_C_ (brown) and D405_C_ (red) are also highlighted with black dashed circles (top and bottom), while the location of N234 glycan is highlighted with orange dashed circles.

We can also characterize the 1Up-to-2Up transition (Figure S22a), where RBD_B_ undergoes the transitions. The increase in RBD_B_ hinge angle is accompanied with a slight decrease in RBD_A_ hinge angle. The latter decrease likely relates with the increases of the contacts and HBs between RBD_A_ and RBD_C_ from 1Ua, the top populated cluster in 1Up, to two-Up-like structures, for instance, K378_A_-E484_C_, Y369_A_-N487_C_ and D427_A_-Y505_C_ (Figures S20 and S21). The numbers of contacts and HBs in two-Up-like structures are much less than those in one-Up, suggesting that a drastic reduction of inter-domain interactions is required toward a full two-Up conformations. No specific protein-glycan interactions are found to support the 1Up-to-2Up transition.

Finally, we examine the effect of glycans on the Down-to-Up transitions. Figure S22b shows the distributions of gREST_Down w/o glycan simulation on the Hinge_A_-Hinge_B_ space. Without glycans on the surface of S-protein, the structure distribution becomes wider, suggesting that S-protein is more flexible without glycans. In addition, Down conformations are more diverse and asymmetric as one RBD has larger hinge angle distribution than the others. The transition mechanisms seem to be the same as those with glycans: RBD_A_ and RBD_C_ form transient interactions to support the Down-to-1Up transition. However, the two-Up-like structure (2Ub_L_) is more connected with Down distribution, suggesting that structural integrity of S-protein is lost without glycans on the surface.

### Searching for cryptic binding pockets in the intermediate structures

We applied a machine learning based algorithm (P2Rank)^45^ to search for the formation of druggable pockets in the intermediates structures, I2a, I3a and I3b (Figure S23a). The same search was also carried out for Down and one-Up structures for comparison. Figure 5a shows the formation of two cryptic pockets (pocket1 and pocket2) at the interface of RBD in I2a, which are not observed in Down or one-Up. Table S4 lists these predicted pockets showing relatively high scores in all the intermediates. To test the druggability of the two pockets, we performed virtual screening of FDA approved drugs from ZINC database^46^, where we docked 2,115 molecules to the RBD interfaces in I2a, I3a and I3b (following the procedure sketched in Figure 23c). Table S5 shows a list of the index (ID) and binding energies of top ranked molecules. Notably for Nilotinib (Figure 5b), which is ranked at the top 5 and 6, the previous study has shown that it affects the SARS-Cov-2 infectivity^47^. The authors attributed the effect to the reduction of cell entry, but the mechanism remains unknown^47^. Figures 5c and S23d show the top-ranked binding poses of Nilotinib in the three intermediates. Nilotinib binds to either pocket1 or pocket2 in all first 9 binding modes with high binding affinity (Table S6). These results serve the two cryptic pockets as potential targets to stabilize intermediate structures and to prevent the formation of one-Up conformation responsible for the viral entry.

**Figure 5.**
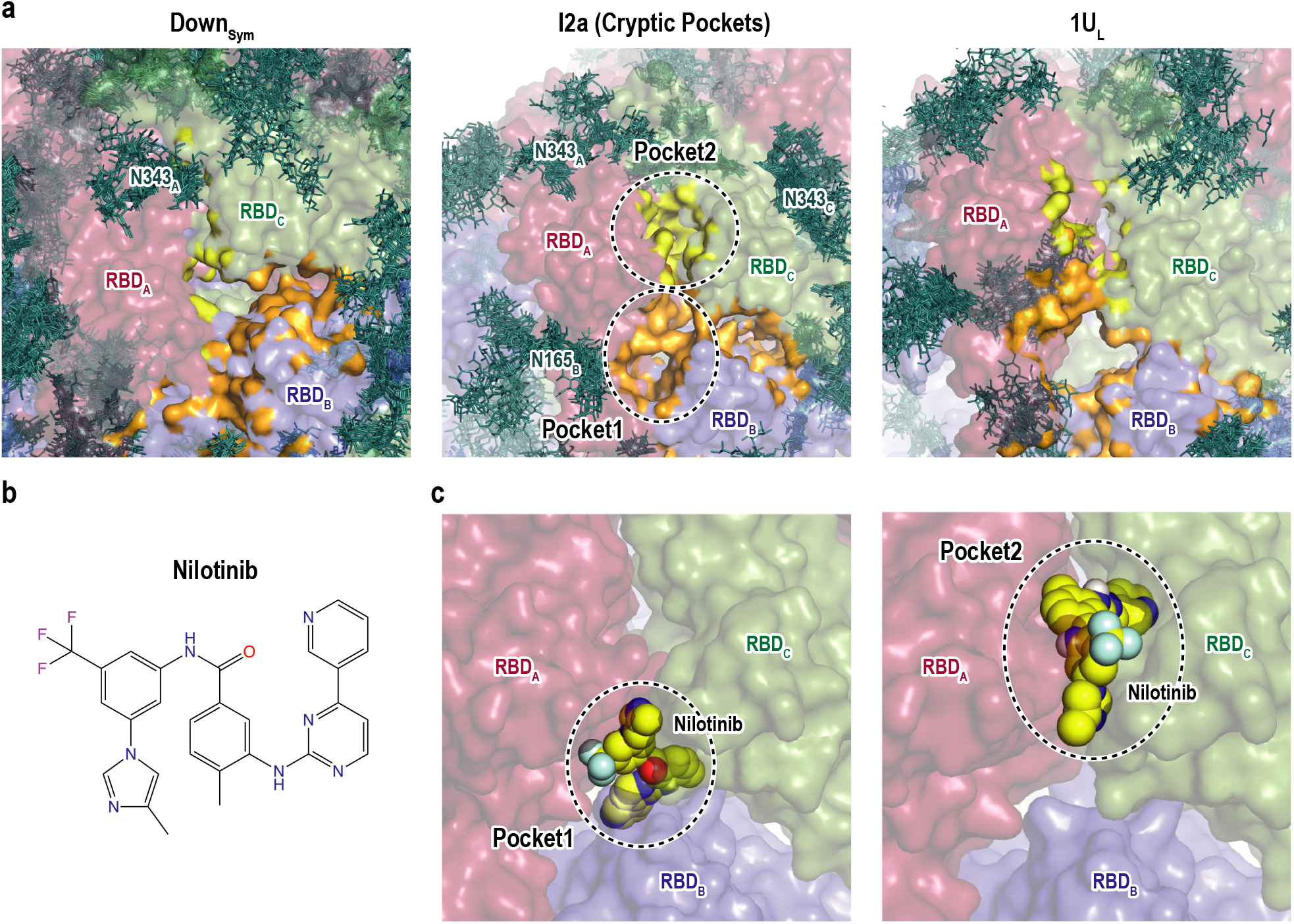
Druggable cryptic pockets. **a)** Snapshots of RBD interface in Down symmetric (Down_Sym_), Intermediate 2a (I2a) and 1Up like (1U_L_) conformations. Chain A, B, and C in the protein are shown in red, blue, and green surfaces, respectively, while a collection of glycans from 10 snapshots are shown in stick representation. The cryptic pockets predicted for I2a using P2Rank software are shown in orange (Pocket1) and yellow (Pocket2), respectively. These pockets disappear in both Down_Sym_ and 1U_L_. **b)** Chemical structure of Nilotinib and **c)** the docked poses (top and third ranked) to two cryptic pockets in I2a by Autodock Vina. The pockets are highlighted with black dashed circles and the nilotinib is shown in sphere representation in yellow.

## Discussion

### Sampling the conformational space of S-protein

Due to the importance of the conformational changes of S-protein in the infection mechanisms and the rational design of antiviral drugs or antibodies, many extensive simulation studies on S-protein have been carried out since the start of the pandemic. The latest challenges involve the use of a million of distributed computer resources, Folding@home, to realize MD simulation of the SARS-CoV-2 proteome for 0.1 second, in total. An alternative approach is to incorporate experimental observation to effectively sample different states. Brotzakis and co-workers recently applied the Cryo-EM Metainference method to determine the opening pathway and intermediates based on the experimental density map^44^. Yet another approach is to use enhanced sampling methods. Sztain and co-workers performed the weighted ensemble simulations, collecting 130 microseconds trajectories, to characterize the opening pathway^35^. Fallon et al. employed the umbrella sampling, nudged elastic band, and steered MD to characterize the transition pathway and associated cryptic pockets^33^. Our gREST_SSCR simulations aimed to enhance sampling of multi-domain proteins like S-protein by exchanging the solute temperatures of selected surface charged residues. It is free from reaction coordinates and bias potentials, which were used in many of the previous enhanced sampling simulations of S-protein. By using Fugaku supercomputer as well as GENESIS program (ver2.0) designed for achieving high scalability on that system^37,38^, the simulations explored a wide conformational space covering Down, one-Up, one-Open, and two-Up-like structures. The structure ensembles that we obtained in the two gREST_SSCR simulations agree with the Cryo-EM structure distributions and smFRET data, suggesting inherent flexibility of S-protein structures even without its ligand, such as hACE2 or nAbs.

Besides the enhancement of conformational sampling by gREST_SSCR, there are several remaining issues in the simulations. Within our computational time, one single simulation cannot cover a whole free-energy landscape containing all the important structures of S-protein. Three-Up structures, which were found in Cryo-EM, for instance, PDB:7DCC^41^, were not observed in our simulations. As we discuss in the next section, glycans on the surface of S-protein seem to have dual roles in the transitions, either helping or hindering the Down-to-Up transitions, which makes conformational sampling via MD simulations more difficult. However, conformational sampling via gREST_SSCR is much better than cMD without any pre-defined reaction coordinates and/or bias potentials, providing sufficient structural information to investigate inherent flexibility of S-protein and the transition mechanisms.

### Role of Flexible Down structures in the Down-to-Up transitions

In this study, we observed inherent flexibility of S-protein structures in absence of its ligand, exploring Down, one-Up, one-Open, and two-Up-like structures at 310 K. The trimeric RBDs in S-protein is intrinsically flexible even in Down form, as their interfaces are electrostatically repulsive as shown in our previous study^31^. Not only symmetric structures but also anti-symmetric structures are also observed in Down, suggesting the existence of “flexible Down structure” (Figure S24). This flexibility can initiate the Down-to-1Up transition, by reducing inter-domain interactions found in Down_Sym_ and allowing the insertion of a glycan at N343_B_ underneath of RBD_A_ (Figure S25). This is consistent with the experimentally mobile RBD conformations in Down observed by Ke et al.^11^ and Wrobel et al.^13^.

Following the flexible Down structure, our simulations elucidate molecular mechanisms underlying the Down-to-1Up transition involving RBD_A_ and the 1Up-to-2Up transitions involving RBD_A_ and RBD_B_. This sequence of events can be explained via the changes of inter-domain interactions as well as protein-glycan interactions. Since RBD_A_-RBD_C_ interaction becomes stronger in the Down-to-1Up transition as we see in Figures 4, S20, and S21, RBD_B_ is more mobile for the next 1Up-to-2Up transition. Among the inter-domain interactions, we pointed out the importance of transient salt-bridges and HBs between R408_A_-D405_C_ and R408_A_-D406C for stabilizing the intermediates (Figures 4 and S21). The salt-bridge between R408_A_ and D405_C_ was also observed in the weighted ensemble simulations reported by Sztain et al.^35^ This scenario is perturbed via protein-glycan interactions, in particular, those involving three glycans at N165_B_, N234_B_, and N343_B_ as pointed out in previous studies^29,30,31^. In the current study, we highlighted the dual roles of these three glycans: the stabilization of one of the states and the driving force toward the other state. The glycans at N165 stabilize Down, representing a barrier for the Down-to-Up transition, while the glycan at N343_B_ drives the transition. Finally, the glycan at N234_B_ stabilizes one-Up-like conformation. This picture coincides with the experimental results by Henderson et al.^48^, where the population of one-Up drastically is reduced by the glycan deletions at N234 but increases by those at N165. As suggested previously^30^, the glycans at N165 may also stabilize one-Up conformation but as a minor role because it can adapt various orientations to that conformation (Figure S25). The role of glycans at N343 is noteworthy in that it helps the position of RBD lift up from Down and supports it throughout the intermediates. This well explains the 20-fold reduction of the infectivity upon N343Q mutation observed by Li et al.^49^. Recently, a similar role of the glycans at N343 has been proposed by Sztain et al.^35^

### Implication for vaccine and drug developments

Since the drug repurposing studies^50,51,52^ so far mostly target Down and Up conformations from Cryo-EM structures, the identification of druggable cryptic pockets in transient intermediates would introduce unprecedented drug targets^44^. Two cryptic pockets at the RBD interface are identified in the highly populated intermediates along the Down-to-1Up transition. From our virtual screening of FDA approved molecules, these pockets are druggable and accommodates several small molecules including Nilotinib. Intriguingly, Cagno et al.^47^ reported that Nilotinib reduces SARS-CoV-2 infection by ~50% via interfering with viral cell fusion and replication. In addition, Nilotinib as well as other top ranked molecules (Table S5) were also predicted to bind RBD in the previous virtual screening studies^52,53^. It is yet unclear if the binding of molecules at RBD interface stabilizes the intermediate states and reduces the population of Up conformation to block ACE2 binding. To shift the conformational equilibrium of S-protein toward the inaccessible Down state for blocking ACE2 and subsequent membrane fusion has been a focus of the recent challenges. Indeed, there have been some reports altering the conformational dynamics of S-protein either via site-specific mutations, disulfide bonds and binding to small molecule^14,25,39,54,55,56^. In this study Nilotinib is only shown as an example, while further studies to identify better molecules targeting the intermediate states would pave the way to hinder viral entry.

The understanding of antibody responses is critical for vaccines and neutralizing antibody developments. The large exposure of RBD_A_ in Up is shown in the current study, which is consistent with several previous studies^22,30,57^. We found that the accessibilities of RBM and the antibody epitopes, including mutational sites, depend sensitively on the RBD conformation, suggesting that each of the Up conformations (1U, 1U_O_ and 2U_L_) tends to bind with distinct classes of antibodies. From our inspection, 1U_O_, which largely exposes RBM but the antibody epitopes of only Class I and II, is the potentially active conformation for both the ACE2 binding and evading from the antibody attack. Class I and II epitope regions include K417, E484, and N501, and their mutations could effectively enhance the infection either by enhancing the ACE2 binding (N501Y)^5^ or reducing the antibody binding (K417N or E484K)^5^. Intriguingly, HB analysis shows that K417 and E484 respectively contribute to stabilize the intermediates I2a/I3b and 2UaL, likely enhancing the population of one-Up conformations. Note that L452 shows a little accessibility (Fig. 3) in contrast with the reported severe mutational effect to an antibody recognition^5^. The similar observation was also reported previously^42^. A possibility is that L452R mutation affects the structural flexibility of S-protein, which consequently gives rise to the enhanced infection. The last point to mention is that the predicted Up conformations are reasonably aligned with the antibody-bound S-protein Cryo-EM structures (Fig. S13). Hence, the antibody bindings of S-protein can be explained based on the conformer selection mechanism^58^. The exploration of conformational diversity of S-protein together with binding free-energy calculations followed by docking simulation would provide valuable structural insights with possible antibody bindings^59^.

## Methods

### gREST_SSCR simulations

The initial structures for S-protein head regions (residue 28-1135) were prepared based on the Cryo-EM structures PDB:6VXX and PDB:6VYB for the Down and Up states, respectively^20^. 18 N-glycans and 1 O-glycan were added per protomer as suggested in previous mass-spectrometry experiments and computational models^23,60^. A full list of included glycans is shown in Figure S2. CHARMM-GUI^61^ were used to prepare the final model including the addition of glycans, ions (0.15 M NaCl) and water molecules. Three gREST_SSCR simulations were performed two from Down in the presence (500 ns) and absence of glycans (150 ns), and one from Up (300 ns). Eight pairs of charged residues per protomer at the RBD interface were selected in the solute region of gREST. The total of number of atoms in solute was 870. All simulations were performed using 16 replicas covering the solute temperature range from 310 to 545 K while maintaining solvent temperature at 310.15 K. All simulations were performed using the new version of GENESIS MD software that was optimized on Fugaku^37,38^. Further detailed information is given in Supporting Information.

### Comparison between Cryo-EM structures and MD simulations using PCA

We used Cryo-EM structures provided by “Spike protein and spike receptors” in Protein Data Bank (http://www.rcsb.org/covid19, deposited date 2020/02/04-2020/12/11, released by 12/30) in this study. Only structures where the number of residues is greater than 700 and the last residue >= 1122 for each protomer were selected in the analysis. 373 protomers of 126 structures and 121 trimeric structures from PDB were satisfied with the criteria and used in this study (Table S1). We adopt a method representing the structure with a 9-beads per protomer as used in the previous work of Henderson et al.^7,39^ Unlike their study, 27 beads consisting of three chains are used here. This coarse-grained model consists of 2 beads for RBD, 3 beads for NTD, 1 bead for SD1 and SD2 each, and 2 beads for the S2 region (CD and S2-b) (see Table S2). Principal component analysis (PCA)^62^ was performed using the selected Cryo-EM structures, after converting to the 27 beads model. The PC vectors are calculated upon fitting all the beads. All simulation trajectories were projected onto the PC1 and PC2 vectors and the potential mean forces (PMFs) were calculated for each simulation. The PMF at 310 K in each gREST_SSCR was obtained using all the trajectories via the Multistate Bennett Acceptance Ratio (MBAR) method^40^. The 373 protomers of the Cryo-EM structures were also used to compare hinge/twist angles.

### Pocket Search and Virtual Screening

The P2Rank software^45^ were used to identify potential druggable pockets in intermediate structures. The cluster centres of 12a, 13a and 13b as well as D1Sym and 1U_L_ were used for pocket search. All top ranked pockets were investigated. Pockets at the RBD interface were selected for further analysis as they exist in all three intermediates but vanish in Down and Up, representing potential cryptic pockets. To check the druggability of these pockets, all FDA approved drugs were downloaded from ZINC database^46^. This includes 2,115 molecules representing 1,379 drug candidates. Open Babel was used to convert PDB to PDBQT. AutoDockTools-1.5.6 was used to prepare RBD receptor^63^. AutoDock Vina were used to dock all 2115 molecules and perform virtual screening^64^.

## Data Availability

The trajectories are available from the corresponding authors upon reasonable request. Models and PDBs of representative structures in each cluster are available at https://covid.molssi.org/.

## Code Availability

The source code of GENESIS (https://www.r-ccs.riken.jp/labs/cbrt/) is distributed under the GNU Lesser General Public License version 3.

## Acknowledgements

This research used computational resources of the supercomputer Fugaku (The evaluation environment in the trial phase) provided by the RIKEN Center for Computational Science. The results obtained on the evaluation environment in the trial phase do not guarantee the performance, power and other attributes of the supercomputer Fugaku at the start of its public use operation. The computer resources of Oakforest-PACS were also provided through HPCI System Research project. (Project ID: hp200153, hp200028). The research was supported in part by MEXT as “FLAGSHIP 2020 project”, “Priority Issue on Post-K computer” (Building Innovative Drug Discovery Infrastructure Through Functional Control of Biomolecular Systems), “Program for Promoting Researches on the Supercomputer Fugaku” (Biomolecular dynamics in a living cell/MD-driven Precision Medicine), and MEXT/KAKENHI (Grant Numbers 19H05645 (to YS), 20K15737 (to H.M.D.), 19K12229 (to SR), and 19K06532 (to TM)) and by RIKEN Pioneering Research Projects (Dynamic Structural Biology/Glycolipidologue Initiative/Biology of Intracellular Environments) (to YS).

## Supplementary Information

Supporting information is available for this paper.

