## Supplementary Tables, Figures and Method for "The Inherent Flexibility of Receptor Binding Domains in SARS-CoV-2 Spike Protein" (from another version of this preprint: version 1, 2021-08-06)

### Supplementary Methods

#### Modelling of Spike protein structure for molecular dynamics simulations

The full-length spike (S) protein is formed of three highly glycosylated protomers, with a 1273 residue each. The trimeric structure is divided into three regions, the head region which is consist of S1 and part of S2 subunits (residue 1-1140), the heptad repeat 2 (HR2)/the transmembrane domain (TM) region (residue 1141-1234) and the cytoplasmic tail region (residue 1235-1273)[1,2]. In this study, truncated structures of S-protein including S1 and part of the S2 subunits (residue 28-1135) were used in the simulations. Wherein the starting structure of the Down and Up conformations were based on the Cryo-EM structures, PDB:6VXX and the PDB:6VYB, respectively[3]. The 6VXX PDB structure includes multiple missing regions at the N-terminal Domain (NTD, residue 70-79, 144-164, 173-185 and 246-262), the Receptor binding Domain (RBD, residue 445-446, 455-461, 469-488 and 502) and the S2 subunit (residue 621-640, 677-688 and 828-853). The 6VYB PDB structure has even more missing regions in the RBD with Up form and the adjacent NTD. Although higher resolution Cryo-EM structures were deposited to the PDB later, only these two structures and PDB:6VSB[1] were available when we started in this study. Due to the large size and the presence of multiple missing regions, several modelling strategies were used to complete the structures. Wherein, residues from 28 to 292 of NTD was modelled based on the SARS-CoV crystal structure (PDB:5X4S at 2.2 Å)[4] using Modeller9.19 software[5]. Then part of the modelled region (residue 28-288) was inserted in the Cryo-EM structure upon fitting the backbone of residues 263-290. Similarly, the crystal structure of the RBD domain (PDB:6LZG at 2.45 Å)[6] (residue 336-515) were inserted in the Cryo-EM structures upon fitting the Cα atoms of residues 336-400. Both modelled NTD and RBD regions shows a good alignment with the resolved regions in the 6VXX and 6VYB Cryo-EM structures, see Figure S1. The VMD program[7] were used to superimpose the modelled regions into the Cryo-EM structure. Finally, the missing regions in the S2 subunit were modelled as loop conformations using the top ranked structure from Modeller9.19[5]. A total of 13 disulfide bonds were included in each protomer including the original 12 disulfide bonds in the Cryo-EM structure and one more in the RBD crystal structure. A comparison of our modelled structure and the more recent high-resolution cryo-EM structure (PDB:6ZGE at 2.6 Å)[2] shows a very good agreement, see Figure

S1. 18 N-glycans and 1 O-glycan were added per protomer as suggested in previous mass-spectrometry experiments and a computational model[8,9]. A full list of included glycans is shown in Figure S2. CHARMM-GUI[10] were used to make the final model including the addition of glycans, ions (0.15 M NaCl) and water molecules. In total, three S-protein models were built including the Down conformation in the absence of glycan, glycosylated S-protein in Down, and the glycosylated S-protein in Up conformation (Figure S1d). The total number of atoms in each model are 657,411, 654,427 and 654,494, respectively, with the average box lengths of 186.947, 186.452 and 186.475Å after equilibration, respectively. Finally, the RBD/SD1 monomer models were made by truncating one protomer from the abovementioned Down and Up models, including residues 315-595 (Figure S8b).

#### Computational methods of gREST_SSCR simulations

We recently proposed an enhance sampling method, the generalized replica-exchange with solute tempering of selected surface charged residue (gREST_SSCR)[11] to enhance large domain motions in multi-domain proteins. In this method the Coulomb and Lennard Jones parameters of surface charged residues at the domain interfaces are selected as a solute region in gREST[12]. In this study to enhance conformational dynamics of S-protein, we performed gREST_SSCR simulations, wherein charged residues at the interfaces between two RBD domains, between RBD and NTD, and between RBD and S2 were selected as the solute region (Figure S3a). In total, 16 residues in each protomer, consisting of 8 positive and 8 negative charged residues, were selected as solute in gREST: K113, K378, K386, R408, K417, K462, R466, R983, E132, E169, D198, D405, E406, D420, D428, and E471. All simulations were performed using 16 replicas covering a solute temperature parameter range from 310.00 to 545.00 K while maintaining solvent temperature at 310.15 K in NVT ensemble. We carried out three gREST_SSCR simulations two from Down in the presence (500 ns) and absence of glycans (150 ns), and one from Up (300) ns.

All simulations were performed using the new version of GENESIS MD software that was optimized on Fugaku[13,14]. The overall performance of gREST_SSCR simulations using 16 replicas is 52 ns/day using 2,048 nodes on Fugaku. CHARMM 36m force field were used for protein (C36m), carbohydrate and ions, while CHARMM TIP3P was used as a water model[15,16]. gREST_SSCR simulations were performed after a series of equilibration steps. First modeled systems were minimized for 10,000 steps, while applying positional restraint on the backbone atoms. Second, using leap-frog integrator and the Langevin thermostat, we heated the simulation systems to 310.15 K in a step wise manner for 100 ps. Third, a series of equilibration steps were performed: 1) MD simulations in the NVT ensemble using the velocity Verlet integrator with stochastic velocity rescaling thermostat[17], 2) those in the NPT ensemble with stochastic velocity rescaling thermostat and MTK barostat[18,19], (note that all previous steps also included a weak restraints on side chain and glycan dihedral angles), 3) after removing all restraints, another MD simulation in the NPT ensemble were performed as equilibration using the same protocol, 4) MD simulation in the NVT ensemble were followed as the second equilibration using the same thermostat and the multiple time-step integrator (MTS) with a fast motion time step of 2.5 fs, and slow motion every 5 fs[19,20]. 5) Prior to production run, a 2 ns equilibration was performed for 16 replicas. Production runs were then performed for 150, 500 and 300 ns per replica in gREST_Down w/o glycan, gREST_Down and gREST_Up simulations, respectively. At every 20 ps, replica exchanges were attempted, and trajectories were saved. Electrostatic interactions were computed by smooth particle mesh Ewald (SPME)[21] method with 128 × 128 × 128 grids and the 6$^{th}$-order B-spline function. Temperature is evaluated using the group-based approach with an optimal temperature evaluation, and thermostat is applied at every 10 steps[22]. Classical MD simulation of RBD/SD1 monomer structures were performed for 300 ns. Two independent simulations were performed starting from Up and one from Down. In all simulations, water molecules were constrained with SETTLE, while bonds involving hydrogens were constrained with SHAKE/RATTLE algorithm[23].

#### Simulation trajectory analysis

To characterize the RBD motions, two main criteria are considered: the Cα atoms root mean square deviations (RMSD) of RBD upon fitting the S2 Cα atoms of Cryo-EM structure (residue 689-827 and 854-1134) and the RBD hinge and twist angles. Hinge and twist angles represent relative domain motions of RBD, wherein the hinge angle describes the Down/Up transition while the twist angle describes RBD side motion. The hinge angle is defined with three points, the center of mass of the Cα atoms in the SD1 core (residue 324-329, 531-538, and 537-590), the top residues of SD1 (residues 328, 329, 530, 531, 543 and 544) and the center of mass in the RBD core (residues 335-466 and 491-526). To define the twist angle, one more point was added at the lower part of RBD (residues 335, 336, 361, 362, 524, and 525). To examine the intra-domain stability, we computed the Cα atom root mean square deviations (RMSD) of RBD (residues 333-528) and NTD (residues 28-306).

The *k*-means algorithm in GENESIS software package were used to classify the conformations of S-protein in MD simulations at 310 K. Hereafter, all the analysis is carried out to obtain the canonical ensembles at 310 K. The number of clusters in *k*-means clustering was set to eight in all cases. The cluster analysis was performed with the same fittings used in the RMSD analysis of RBD. Only the Cα atoms included in the original Cryo-EM structures (PDB:6VXX and PDB:6VYB) were included to avoid flexible regions in our analysis. Furthermore, the distributions of hinge and twist angles for all the 8 clusters were calculated for each protomer (in total, 6 angles) and subsequently the number of clusters were increased until the hinge/twist distribution showed the minimal overlaps (Figures S14-S16). In this procedure, 12, 13 and 13 clusters were obtained in gREST_Down w/o glycan, gREST_Down and gRSET_Up simulations, respectively (Table S3).

Due to the homo-trimeric nature of S-protein, protomers are indistinguishable in the structure. gREST_SSCR enhanced motions of RBD regions so that we don't know which protomer reveals large-scale conformational motions in any replicas. For instance, Figure S6b shows that $RBD_A$ undergoes large transition in replica 1, while $RBD_C$ shows large motion in replica 16. To clarify the discussion in this paper, we applied a rotational scheme that makes $RBD_A$ undergo the largest conformational transitions in the following ways: 1) We identify all replicas that show significant RBD motions with a hinge angle > 130° in $RBD_B$ or $RBD_C$. 2) We rotate the conformations of those selected replicas where $RBD_B$ or $RBD_C$ becomes $RBD_A$ while rotating the rest of the molecule including glycans (Figure S6a). 3) We confirm the rotation scheme by comparing hinge/twist angle free energy maps before and after rotation (Figures S6c, S6d, S7a, and S7b). We also compared principal component analysis (PCA) before and after rotations. 4) In cases of two RBDs showing large hinge angles in the same replica, the protomer with the highest RBD hinge angle becomes $Chain_A$.

Hydrogen bonding (HB) and contact analysis were also performed for major clusters, wherein a 75% and 50% probability threshold were used for the heavy atoms contacts and HB residue pairs selection in Figures S20 and S21, respectively. The correspondence analysis to the previous smFRET experiment[24] was performed upon calculating the COM distance from residues 425-431 to residues 554-561 using the Cα atoms (Figure S17a). Experimental statistical ratio[24] of 77 and 23 % was used to combine gREST_Down and gREST_Up simulation results, respectively as shown in Figure S17d. The VMD and PyMOL programs were used for trajectory and structure visualization[7,25].

Solvent Accessible Surface Area (SASA) values were calculated using the measure SASA function in VMD[7]. The restrict option, which considers only solvent accessible points near the user specified region, was used for per-domain and per-residue SASA calculations. A range of probe radius, including 1.4 Å (a sphere of water) and 7.2 Å (approximating the hypervariable loops of antip-gp120 antibodies)[26], was used for SASA calculations. SASA values were calculated for different RBD conformations, Down, 1U, $1U_O$, and $2U_L$. Down represent the sum of $Down_{Sym}$ and $Down_{Asym}$, the rest is defined in Table S3. For each conformation, 30 snapshots close to the cluster center were extracted and used for analysis. The calculated SASA values were mapped on the structure using PyMOL software[25]. For comparison, we confirmed that our SASA calculations give the results in consistent with the previous work by Amaro and co-workers (Figure S9)[27].

**Supplementary Table 1: Cryo-EM structures used in the PCA analysis.**

"Form" is determined by the number of chains which take Down form of RBD in S-protein (3: Down, 2: 1UP, 1: 2UP, 0: 3Up, respectively). "Different protein" means whether there are protein chain(s) other than S-protein (uniprot: P0DTC2) in Cryo-EM or X-ray structures.

| PDB | Form | Different protein | PDB | Form | Different protein | PDB | Form | Different protein |
|---|---|---|---|---|---|---|---|---|
| 6VXX | Down | No | 7K8S | Down | Yes | 7A25 | 1UP | Yes |
| 6WPS | Down | Yes | 7K90 | Down | Yes | 7A94 | 1UP | Yes |
| 6X29 | Down | No | 7KDG | Down | No | 7AD1 | 1UP | No |
| 6X2C | Down | No | 7KDI | Down | No | 7BYR | 1UP | Yes |
| 6X6P | Down | No | 7KDK | Down | No | 7CHH | 1UP | Yes |
| 6X79 | Down | No | 7KE4 | Down | No | 7CN9 | 1UP | No |
| 6XEY | Down | Yes | 7KE6 | Down | No | 7CWM | 1UP | Yes |
| 6XF5 | Down | No | 7KE7 | Down | No | 7DD8 | 1UP | Yes |
| 6XLU | Down | No | 7KE8 | Down | No | 7DDN | 1UP | No |
| 6XM5 | Down | No | 7KKK | Down | Yes | 7DF4 | 1UP | Yes |
| 6ZB4 | Down | No | 7KKL | Down | Yes | 7DK5 | 1UP | Yes |
| 6ZB5 | Down | No | 7L02 | Down | Yes | 7JV4 | 1UP | Yes |
| 6ZGE | Down | No | 7L06 | Down | Yes | 7K8T | 1UP | Yes |
| 6ZGI | Down | No | 7L09 | Down | Yes | 7K8V | 1UP | Yes |
| 6ZOX | Down | No | 6VSB | 1UP | No | 7K8W | 1UP | Yes |
| 6ZOY | Down | No | 6VYB | 1UP | No | 7K8X | 1UP | Yes |
| 6ZOZ | Down | No | 6WPT | 1UP | Yes | 7K8Z | 1UP | Yes |
| 6ZP0 | Down | No | 6X2A | 1UP | No | 7KDH | 1UP | No |
| 6ZP1 | Down | No | 6XF6 | 1UP | No | 7KDJ | 1UP | No |
| 6ZP2 | Down | No | 6XKL | 1UP | No | 7KDL | 1UP | No |
| 7A4N | Down | No | 6XM0 | 1UP | No | 7KE9 | 1UP | No |
| 7CAB | Down | No | 6XM3 | 1UP | No | 7KEA | 1UP | No |
| 7DDD | Down | No | 6XM4 | 1UP | No | 7KEB | 1UP | No |
| 7DF3 | Down | No | 6Z43 | 1UP | Yes | 7KEC | 1UP | No |
| 7JJI | Down | No | 6Z97 | 1UP | No | 7KJ2 | 1UP | Yes |
| 7JV6 | Down | Yes | 6ZGG | 1UP | No | 7KJ5 | 1UP | No |
| 7JWY | Down | No | 6ZHD | 1UP | Yes | 7KNB | 1UP | Yes |
| 6XCM | 2UP | Yes | 6ZXN | 1UP | Yes | 7KNE | 1UP | Yes |
| 7A29 | 2UP | Yes | 7K8Y | 2UP | Yes | 6X2B | 2UP | No |
| 7A93 | 2UP | No | 7KJ3 | 2UP | Yes | 7DCX | 3UP | Yes |
| 7A95 | 2UP | Yes | 7KL9 | 2UP | Yes | 7DK7 | 3UP | Yes |
| 7A96 | 2UP | Yes | 7KMZ | 2UP | Yes | 7JVC | 3UP | Yes |
| 7A97 | 2UP | Yes | 7KNH | 2UP | Yes | 7JW0 | 3UP | Yes |
| 7CAI | 2UP | Yes | 6XCN | 3UP | Yes | 7K4N | 3UP | Yes |
| 7DD2 | 2UP | Yes | 6ZDH | 3UP | Yes | 7KJ4 | 3UP | Yes |
| 7DK4 | 2UP | Yes | 7A98 | 3UP | Yes | 7KMS | 3UP | Yes |
| 7DK6 | 2UP | Yes | 7CAK | 3UP | Yes | 7KNI | 3UP | Yes |
| 7JWB | 2UP | Yes | 7CT5 | 3UP | Yes | 6ZGH | 2 chains | No |
| 7JZL | 2UP | Yes | 7CWN | 3UP | Yes | 6ZOW | 2 chains | No |
| 7JZN | 2UP | Yes | 7CWS | 3UP | Yes | 6ZP5 | 2 chains | No |
| 7K8U | 2UP | Yes | 7CWU | 3UP | Yes | 6ZP7 | 2 chains | No |
| 7K43 | Down | Yes | 7DCC | 3UP | Yes | 7DK3 | 2 chains | No |

**Supplementary Table 2: Definition of protomer coarse-grained particles representing rigid domains for PCA.**

| Rigid domains | Residue numbers |
|---|---|
| NTD | 27-43, 54-271 |
| NTD-b | 116-129, 169-172 |
| RBD | 330-443, 503-528 |
| RBD-h | 403-410 |
| NTD’ | 44-53,272-293 |
| SD1 | 323-329, 529-590 |
| SD2 | 294-322, 591-696 |
| S2-b | 717-727, 1047-1071 |
| CD | 711-716, 1072-1122 |

**Supplementary Table 3: List of clusters for gREST_Down, gREST_Up and gREST_Down w/o glycan simulations.**

| Initial clusters | Refined clusters | Cluster name | Macro clusters |
|---|---|---|---|
| **gREST_Down** | | | |
| C1 | C1 | $D1_{asym}$ | $Down_{Like}$ ($Down_{Asym}$) |
| C2 | C2 | I1a | $Down_{Like}$ (Int1) |
| C3 | C3 | $D1_{Sym}$ | $Down_{Sym}$ |
| C4 | C4(1) | I2a | Int2 |
| C5 | C5(1) | I1b | $Down_{Like}$ (Int1) |
| C6 | C6 | I3b | Int3 |
| C7 | C7(1) | $D2_{asym}$ | $Down_{Like}$ ($Down_{Asym}$) |
| C8 | C8 | $D2_{Sym}$ | $Down_{Sym}$ |
| | C4(2) | I3a | Int3 |
| | C5(2) | I1c | $Down_{Like}$ (Int1) |
| | C7(2) | I2b | Int2 |
| | C7(3) | I2c | Int2 |
| | C7(4) | $1U_L$ | 1Up |
| **gREST_Up** | | | |
| C1 | C1 | $1U_O$ | 1Up |
| C2 | C2 | 1Ub | 1Up (1U) |
| C3 | C3(1) | 1Ue | 1Up (1U) |
| C4 | C4(1) | 1Uc | 1Up (1U) |
| C5 | C5(1) | $2Ua_L$ | 1Up ($2U_L$) |
| C6 | C6 | 1Uf | 1Up (1U) |
| C7 | C7(1) | 1Uh | 1Up (1U) |
| C8 | C8 | 1Ua | 1Up (1U) |
| | C3(2) | 1Ud | 1Up (1U) |
| | C4(2) | 1Ug | 1Up (1U) |
| | C5(2) | $2Ub_L$ | 1Up ($2U_L$) |
| | C7(2) | 1Uj | 1Up (1U) |
| | C7(3) | 1Ui | 1Up (1U) |
| **gREST_Down w/o glycan** | | | |
| C1 | C1(1) | IUa | |
| C2 | C2 | I1b | |
| C3 | C3(1) | I2a | |
| C4 | C4(1) | D3 | |
| C5 | C5 | D1 | |
| C6 | C6 | I1a | |
| C7 | C7 | D4 | |
| C8 | C8(1) | D2 | |
| | C1(2) | 1Ub | |
| | C3(2) | I2b | |
| | C4(2) | 2UIb | |
| | C8(2) | 2UIa | |

**Supplementary Table 4: The RBD interface cryptic pockets predicted by P2Rank[28].**

| Intermediate | Rank | Score | Probability | Residue ids |
|---|---|---|---|---|
| **I2a** | 3 (Pocket_1) | 113.9 | 0.994 | A_403 A_405 A_406 A_408 A_409 A_414 A_415 A_416 A_417 A_420 A_421 A_424 A_455 A_456 A_460 A_473 A_474 A_475 A_476 A_477 A_478 A_479 A_489 B_335 B_336 B_338 B_339 B_342 B_344 B_345 B_367 B_368 B_372 B_373 B_374 B_375 B_376 B_378 B_380 B_403 B_404 B_405 B_406 B_407 B_408 B_409 B_410 B_411 B_414 B_415 B_416 B_417 B_420 B_421 B_433 B_435 B_436 B_437 B_438 B_439 B_440 B_441 B_449 B_453 B_455 B_493 B_494 B_495 B_496 B_498 B_499 B_500 B_501 B_502 B_503 B_504 B_505 B_506 B_508 B_509 C_368 C_369 C_371 C_372 C_374 C_375 C_376 C_377 C_378 C_379 C_380 C_382 C_384 C_403 C_404 C_405 C_407 C_408 C_411 C_414 C_437 C_501 C_502 C_503 C_504 C_505 C_506 C_508 |
| | 8 (Pocket_2) | 37.0 | 0.896 | A_375 A_376 A_378 A_380 A_404 A_405 A_407 A_408 A_411 A_412 A_414 A_503 A_504 A_508 C_403 C_406 C_409 C_416 C_417 C_418 C_453 C_455 C_493 C_494 C_495 C_496 C_498 C_500 C_501 C_505 |
| **I3a** | 8 (Pocket_1) | 49.6 | 0.942 | A_403 A_405 A_406 A_408 A_409 A_413 A_414 A_415 A_416 A_417 A_418 A_420 A_421 A_424 A_427 A_453 A_455 A_456 A_459 A_460 A_461 A_463 A_505 B_375 B_376 B_377 B_403 B_404 B_405 B_406 B_407 B_408 B_500 B_501 B_502 B_503 B_504 B_505 B_508 C_375 C_376 C_404 C_405 C_408 C_502 C_503 C_504 C_505 C_508 |
| | 9 (Pocket_2) | 38.7 | 0.908 | A_375 A_376 A_378 A_380 A_404 A_405 A_407 A_408 A_411 A_412 A_414 A_502 A_503 A_504 A_505 A_508 C_403 C_406 C_409 C_415 C_416 C_417 C_418 C_449 C_453 C_455 C_493 C_494 C_495 C_496 C_498 C_500 C_501 C_505 |
| **I3b** | 5 (Pocket_2) | 37.8 | 0.904 | A_371 A_374 A_375 A_376 A_377 A_378 A_379 A_380 A_381 A_404 A_405 A_407 A_408 A_409 A_411 A_412 A_414 A_433 A_437 A_439 A_499 A_501 A_502 A_503 A_506 A_508 C_403 C_405 C_406 C_408 C_409 C_414 C_417 C_453 C_455 C_493 C_494 C_495 C_496 C_498 C_500 C_501 C_503 C_504 C_505 |
| | 12 (Pocket_1) | 26.2 | 0.815 | A_420 A_421 A_455 A_456 A_457 A_460 A_473 A_475 A_476 A_487 A_489 B_375 B_376 B_378 B_407 B_408 B_433 B_437 B_438 B_439 B_441 B_502 B_504 B_505 B_506 B_508 |
| | 16 | 21.6 | 0.747 | B_415 B_416 B_417 B_420 B_421 B_455 B_456 B_457 B_458 B_460 B_473 B_474 B_475 B_476 B_477 B_478 B_487 B_489 B_490 B_493 C_367 C_369 C_370 C_371 C_372 C_373 C_374 C_375 C_376 C_377 C_384 |

**Supplementary Table 5: List of the top ranked molecules from the virtual screening of 2115 FDA approved drugs to RBD interface in I2a, I3a and I3b intermediate structures.**

| Rank | ZINC ID | I2a* | I3a* | I3b* |
|---|---|---|---|---|
| 1 | ZINC000001612996 | −11.9 | −9.5 | −9.4 |
| 2 | ZINC000052955754 | −11.3 | −9.1 | −9.3 |
| 3 | ZINC000003978005 | −11.1 | −9.9 | −9.4 |
| 4 | ZINC000169289767 | −11.1 | −10.0 | −10.1 |
| 5 | ZINC000006716957 | −10.9 | −9.6 | −9.5 |
| 6 | ZINC000006716957 | −10.8 | −10.0 | −9.4 |
| 7 | ZINC000003978005 | −10.8 | −9.8 | −9.8 |
| 8 | ZINC000003932831 | −10.8 | −9.3 | −9.1 |
| 9 | ZINC000003978005 | −10.7 | −10.4 | −10.4 |
| 10 | ZINC000052955754 | −10.6 | −9.8 | −9.7 |
| 11 | ZINC000052955754 | −10.5 | −9.7 | −9.7 |
| 12 | ZINC000053683151 | −10.4 | −9.8 | −9.0 |
| 13 | ZINC000064033452 | −10.4 | −9.9 | −9.3 |
| 14 | ZINC000011679756 | −10.3 | −9.2 | −10.0 |
| 15 | ZINC000036701290 | −10.3 | −9.6 | −9.5 |
| 16 | ZINC000084668739 | −10.2 | −10.1 | −9.3 |
| 17 | ZINC000003927822 | −10.1 | −9.0 | −9.4 |
| 19 | ZINC000164528615 | −10.1 | −9.6 | −9.0 |
| 20 | ZINC000100378061 | −10.0 | −9.8 | −9.0 |

Molecules are ranked based on binding energy to I2a while binding energy to other intermediates (I3a and I3b) are also shown.
* Binding energy in kcal $mol^{-1}$.
Rank 5 and 6, shown in red, represent Nilotinib.

**Supplementary Table 6: Nilotinib binding energy to I2a, I3a and I3b intermediates.**

| | **I2a** | **I3a** | **I3b** |
|---|---|---|---|
| **Mode** | **Affinity (kcal $mol^{-1}$)** | | |
| **1** | −10.9 | −9.6 | −9.5 |
| **2** | −10.9 | −9.6 | −9.4 |
| **3** | −10.8 | −9.5 | −9.3 |
| **4** | −10.8 | −9.4 | −9.2 |
| **5** | −10.6 | −9.3 | −9.2 |
| **6** | −10.5 | −9.2 | −9.2 |
| **7** | −10.4 | −9.0 | −9.2 |
| **8** | −10.4 | −8.8 | −9.1 |
| **9** | −10.3 | −8.7 | −9.1 |

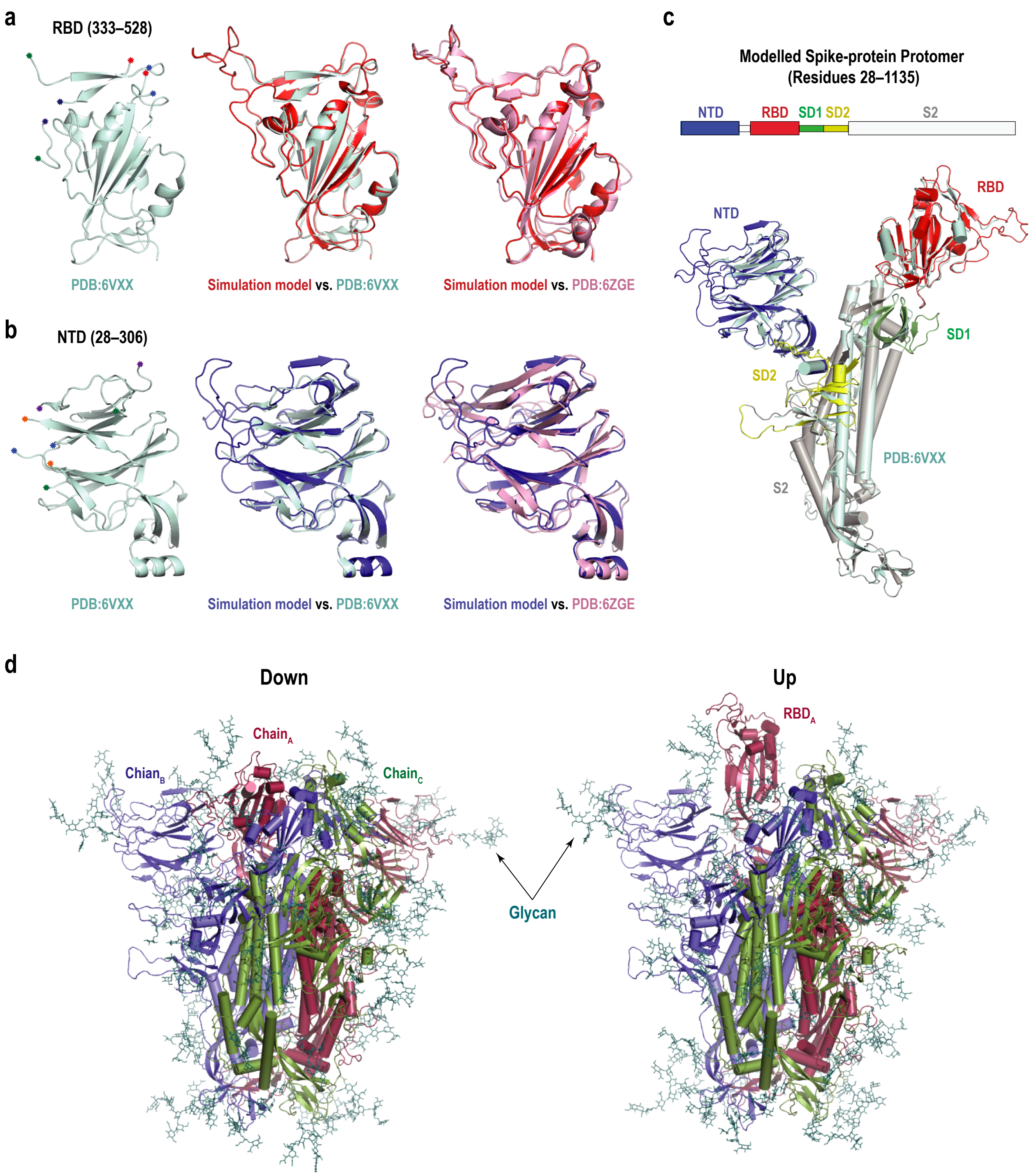


**Figure S1. Structural model of the spike protein used in this study. a)** Left: cartoon representation of the RBD in the original cryo-EM structure (PDB:6VXX). The terminal residues before and after the missing loops are highlighted with colored dots. Middle: superimposition of our simulation model, which was modeled from PDB:6LZG (red), to PDB:6VXX (cyan). Right: comparison between our simulation model (red) and more recent high-resolution cryo-EM structure in Down conformation (PDB:6ZGE) (pink). **b)** Left: cartoon representation of the NTD in the original cryo-EM structure (PDB:6VXX). The terminal residues before and after the missing loops are highlighted with colored dots. Middle: superposition of our simulation model (blue) to PDB:6VXX (cyan). Right: comparison between our simulation model (blue) and PDB:6ZGE (pink). **c)** Superposition of the modeled protomer (residues 28–1135) to PDB:6VXX (cyan). Besides missing residues in the RBD and NTD, three other regions were also modelled using Modeller[5]. The head of the protomer is composed of S1 and S2 subunits. S1 includes NTD (blue), RBD (red), SD1 (green), and SD2 (yellow). Part of S2 that was included in the simulation model (grey). **d)** Full models used in the simulation of Down and 1Up conformations, where 19 glycans per protomer were added. The protomers are shown in red, blue and green cartoon for chain A, B and C, respectively, and glycans are shown as deep teal sticks.

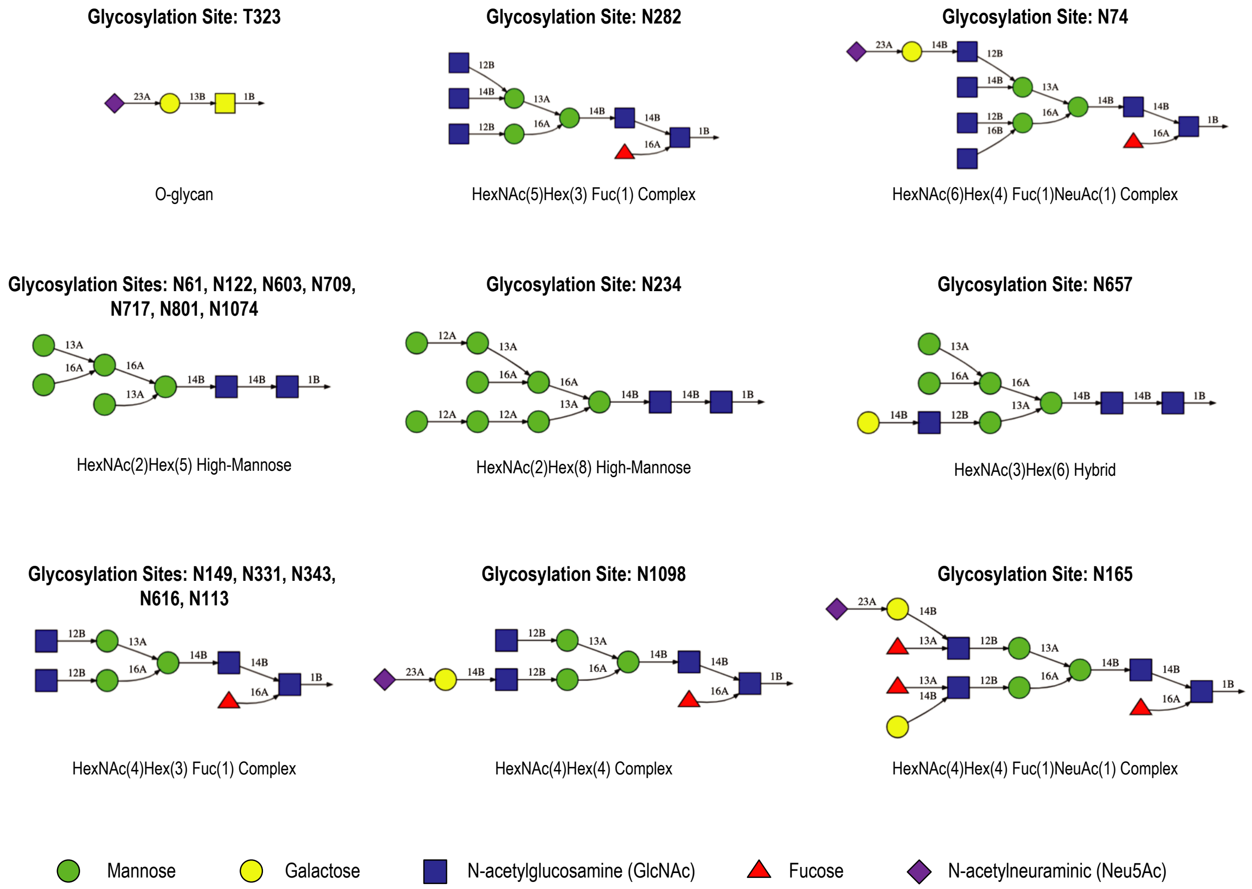


**Figure S2. Glycans in our Spike protein models.** Schematic representations of the glycan structures and types used in gREST_Down and gREST_Up simulations including the location of the glycosylation sites.

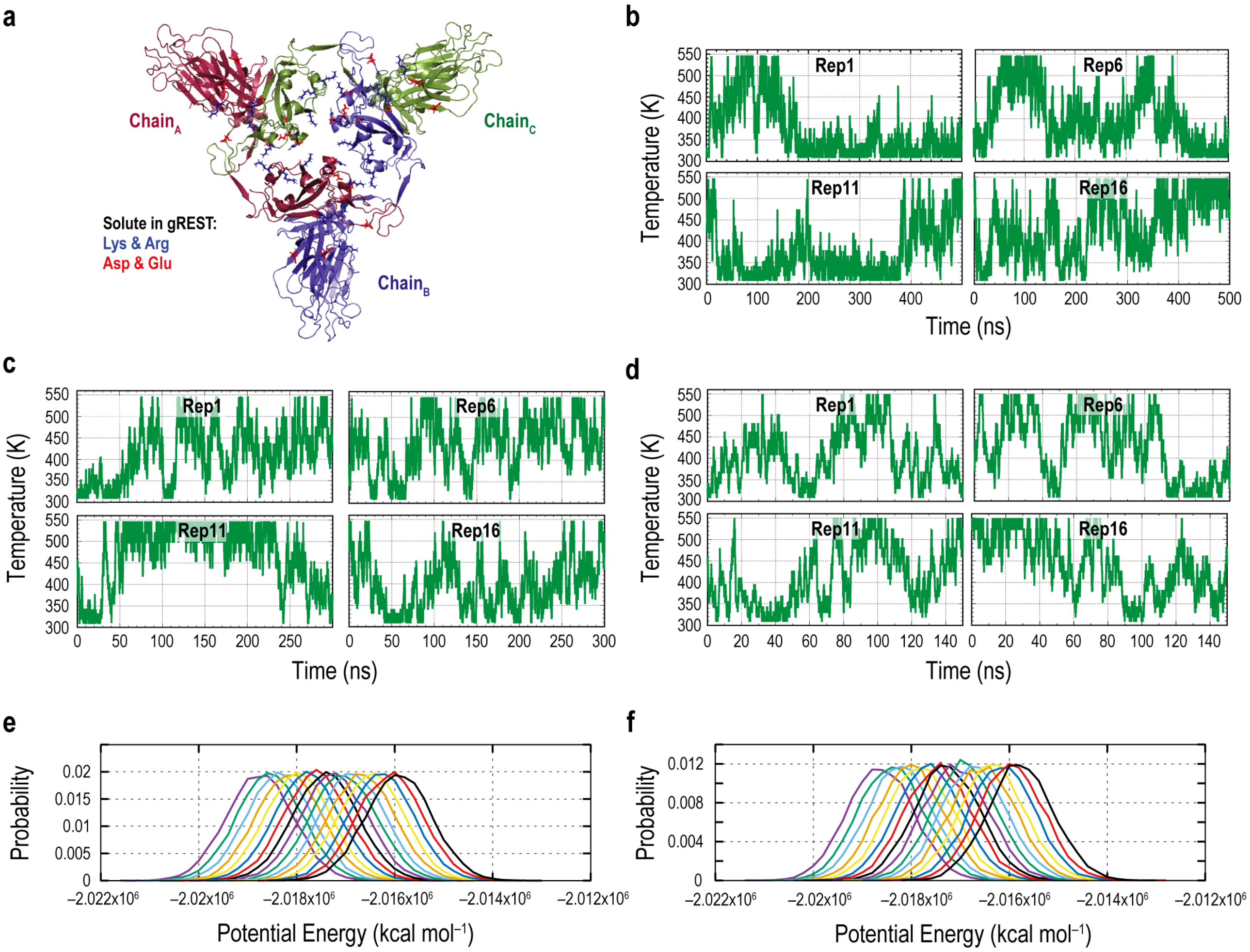


**Figure S3. Performance of the gREST_SSCR simulations. a)** Top view the S1 subunit, where the positively and negatively charged residues used for solute region in gREST are shown as blue and red sticks, respectively. **b, c, d)** Time courses of the temperature in the selected replicas (1, 6, 11 and 16) in the 500 ns gREST_Down (b), 300 ns gREST_Up (c), and 150 ns gREST_Down w/o glycan simulations (d). All three simulations showed random walk in the temperature space. **e, f)** Probability distribution of the potential energies of the sorted temperatures in gREST_Down (e) and gREST_Up simulations (f), both of which showed sufficient overlaps between temperature parameters.

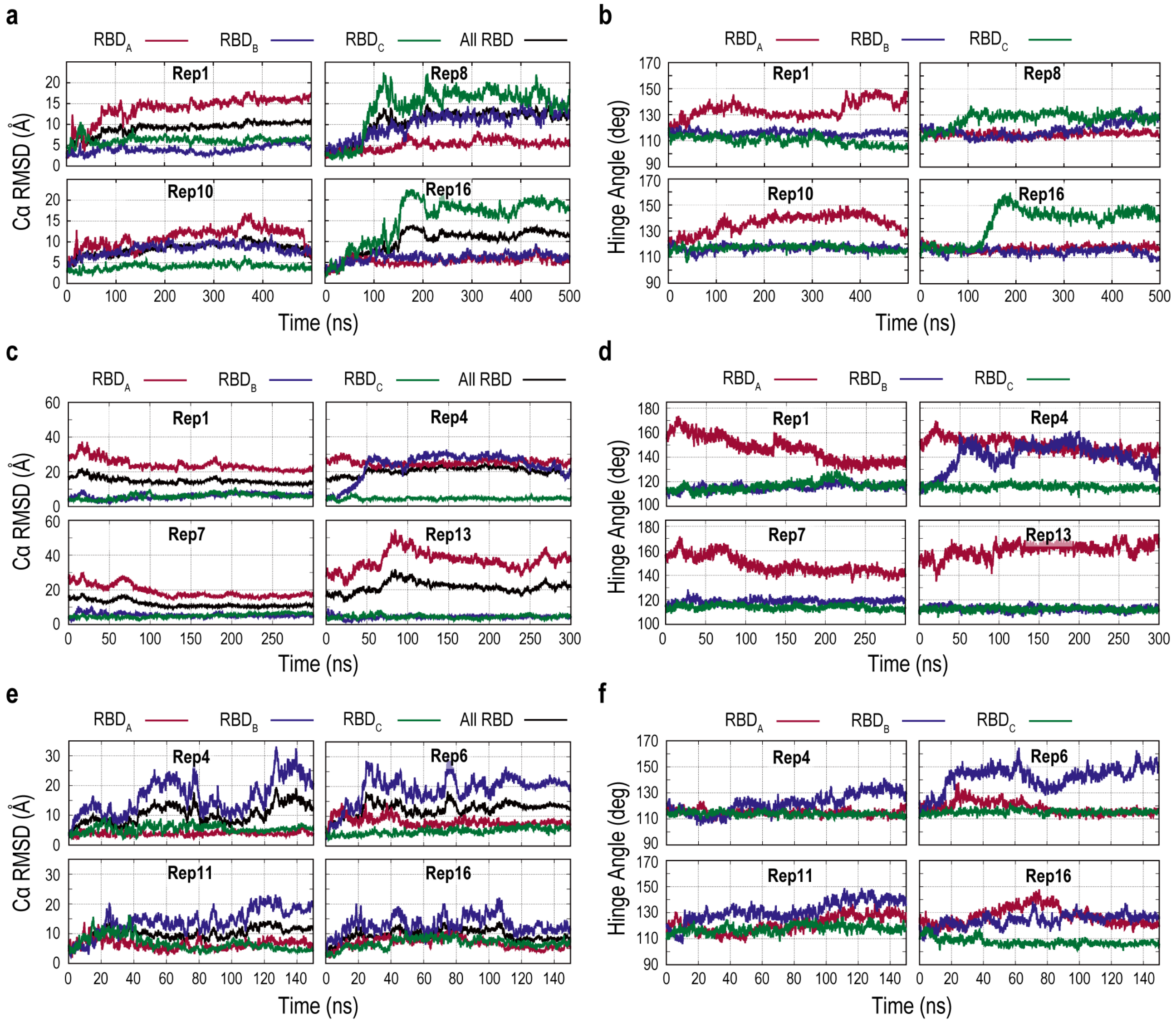


**Figure S4. Characterization of the RBD conformational change in the three gREST_SSCR simulations.** **a, c, e)** Time courses of the root-mean-square deviation (RMSD) of the Cα atoms with respect to Down structure upon fitting the Cα atoms of the S2 subunit in the selected replicas from the gREST_Down (a), gREST_Up (c), and gREST_Down w/o glycan simulations (e). RMSD of the individual RBDs (red, blue, and green for chain A, B, and C, respectively) as well as all three RBDs (black) are shown. **b, d, f)** Time courses of the Hinge angle of RBDs in the gREST_Down (b), gREST_Up (d), and gREST_Down w/o glycan simulations (f).

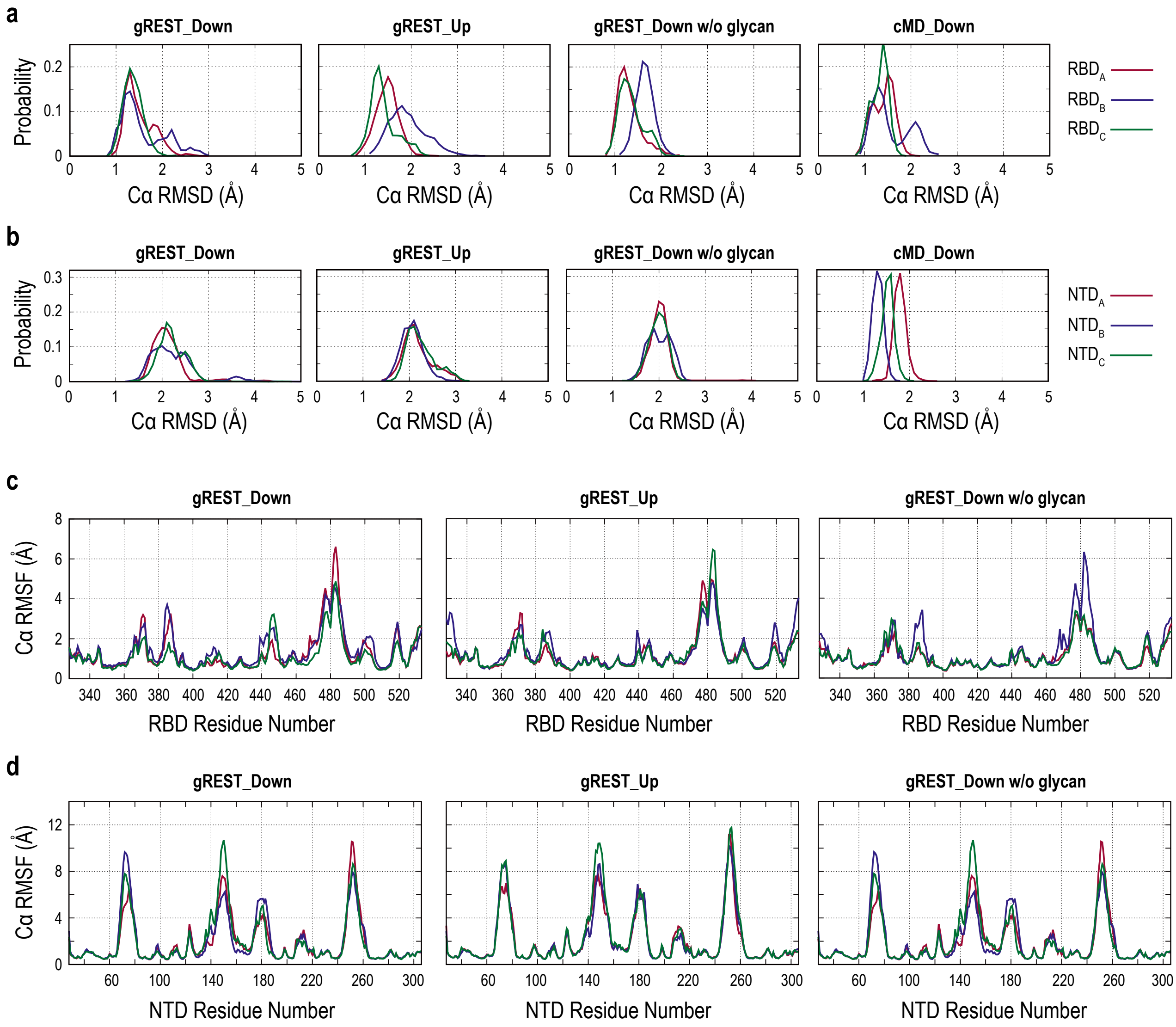


**Figure S5. Analysis of the intra-domain stability of RBD and NTD in the gREST_SSCR simulations. a, b)** Probability distribution of the Cα RMSD of the RBDs (a) and NTDs (b) at 310 K of the gREST_Down, gREST_Up, and gREST_Down w/o glycan simulations as well as our previous 1 μs cMD simulation of the Down conformation[29]. **c, d)** Root-mean-square fluctuation (RMSF) of the Cα atoms of the RBDs (c) and NTDs (d) in the gREST_Down, gREST_Up, and gREST_Down w/o glycan simulations.

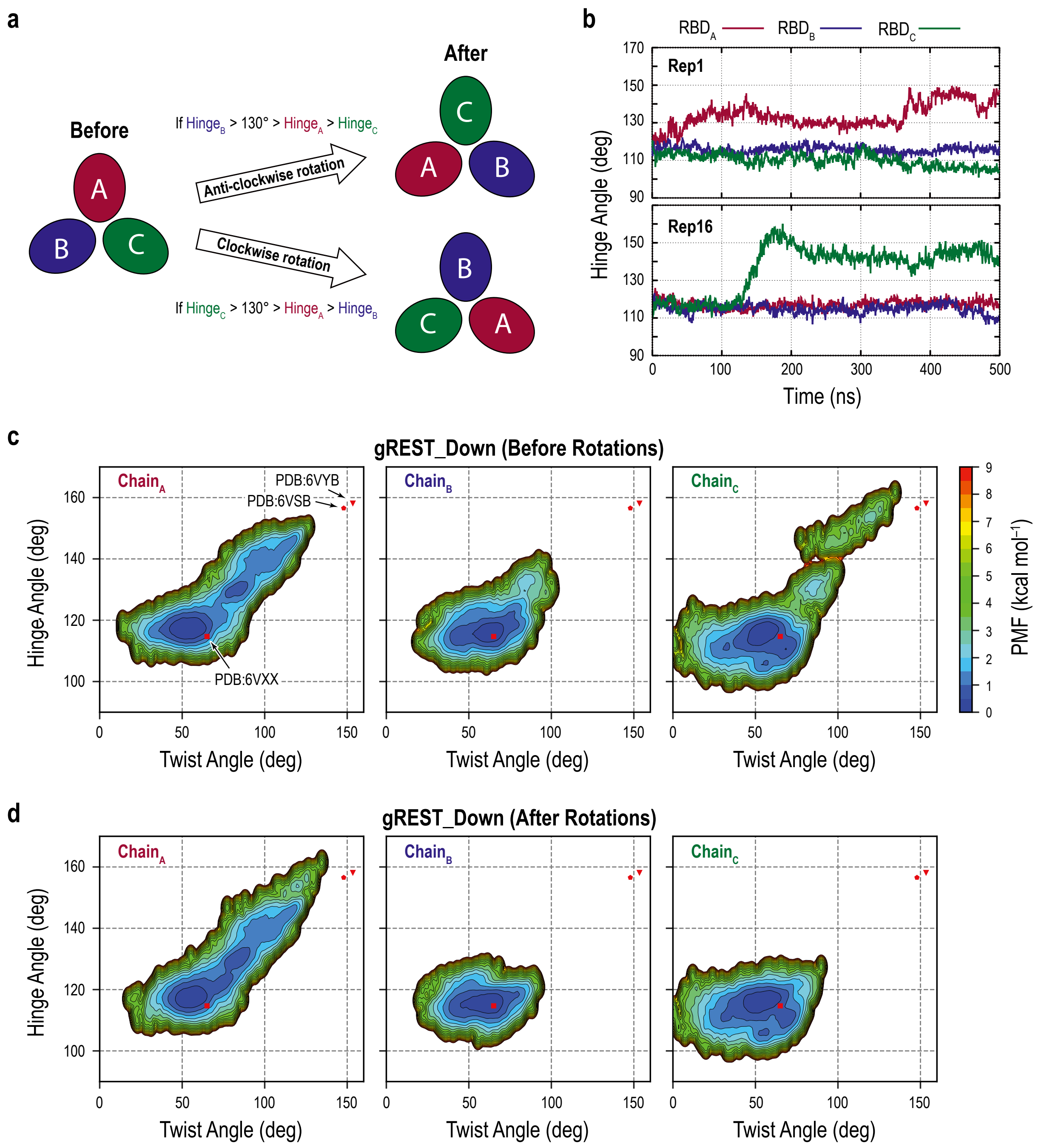


**Fig. S6: Scheme of the protomer rotation, and analysis of the rotated trajectories of the gREST_Down simulation. a)** Schematic representation of the rotation scheme and criteria that was used to define structural changes of RBDs. First, Hinge angle of each RBD is calculated. Then, if the Hinge angle of $RBD_B$ is larger than 130° and also larger than those of $RBD_A$ and $RBD_C$, all three chains are rotated anticlockwise, where the chain index is changed from A to C, B to A, or C to B. Similarly, if $RBD_C$ has the Hinge angle more than 130° and its larger than Hinge angles of $RBD_A$ and $RBD_B$, all chains are rotated clockwise. If $RBD_A$ originally has the largest Hinge angle, no rotation is applied. Rotation was applied to all frames of the selected replicas whose the majority of its snapshots had the above criteria. **b)** Time courses of the Hinge angle in Replicas 1 (upper panel) and 16 (lower panel) from the gREST_Down simulation, where $RBD_A$ and $RBD_C$ showed the largest Hinge angle, respectively. This shows the indistinguishability of RBD in our simulation and the need for rotation scheme for further analysis. **c, d)** Free energy landscape along the Hinge and Twist angles of the three chains before (c) and after the protomer rotation (d). Without the rotation scheme (c), Up like conformations are observed in both chain A and chain C due to the indistinguishability of RBD.

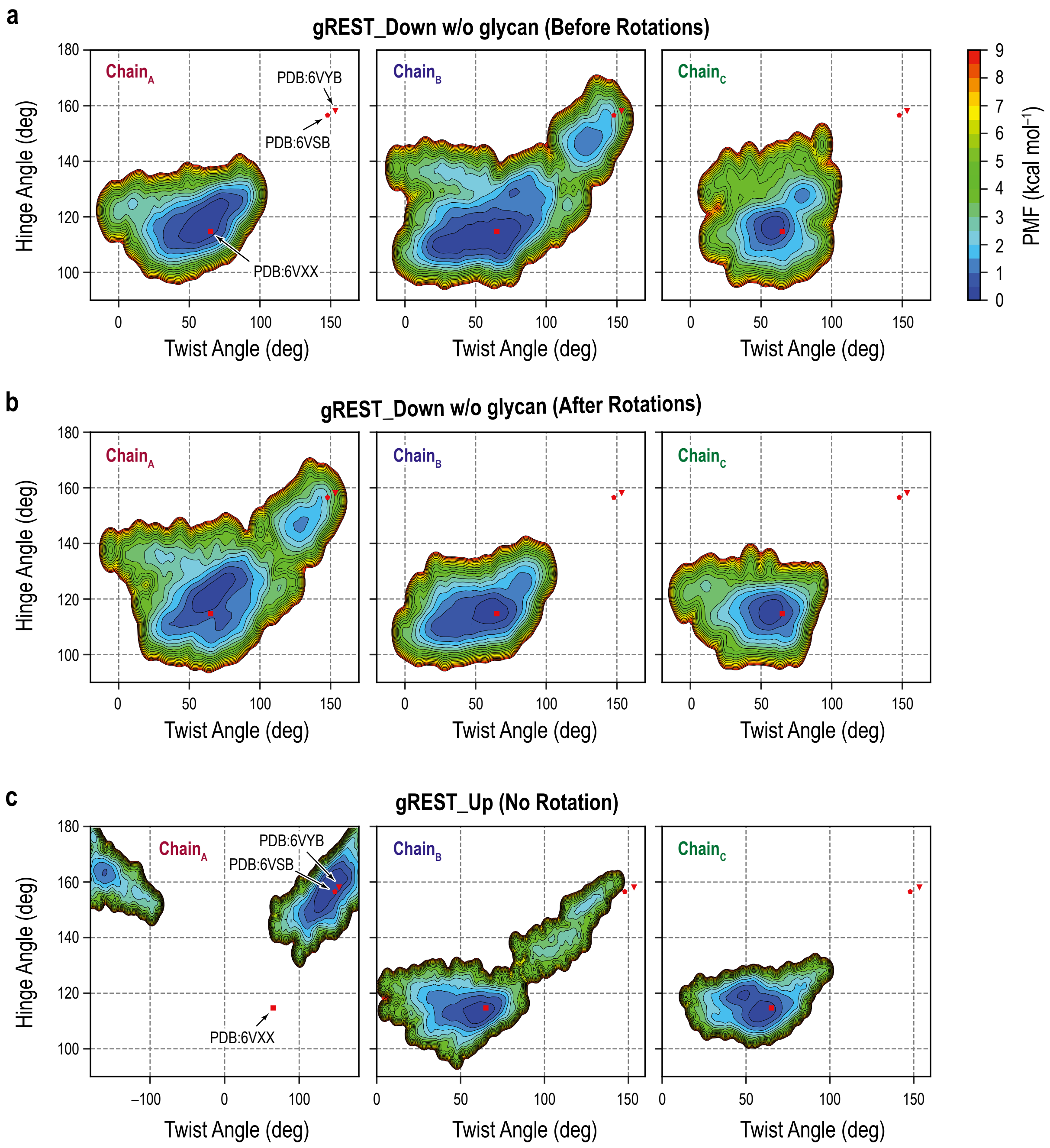


**Fig. S7: Free energy landscape (FEL) along the Hinge/Twist angles in the gREST_Down w/o glycan and gREST_Up simulations. a, b)** FEL in chains A, B, and C before (a) and after the rotation (b) in the gREST_Down w/o glycan simulation. The FELs show significant RBD changes in chain B (or A) and to less extent in chain C. **c)** FEL in chains A, B, and C without rotation in the gREST_Up simulation. The FELs show very large Twist angles in $RBD_A$ as well as the formation of Up conformation in $RBD_B$, demonstrating the formation of 2Up like conformations.

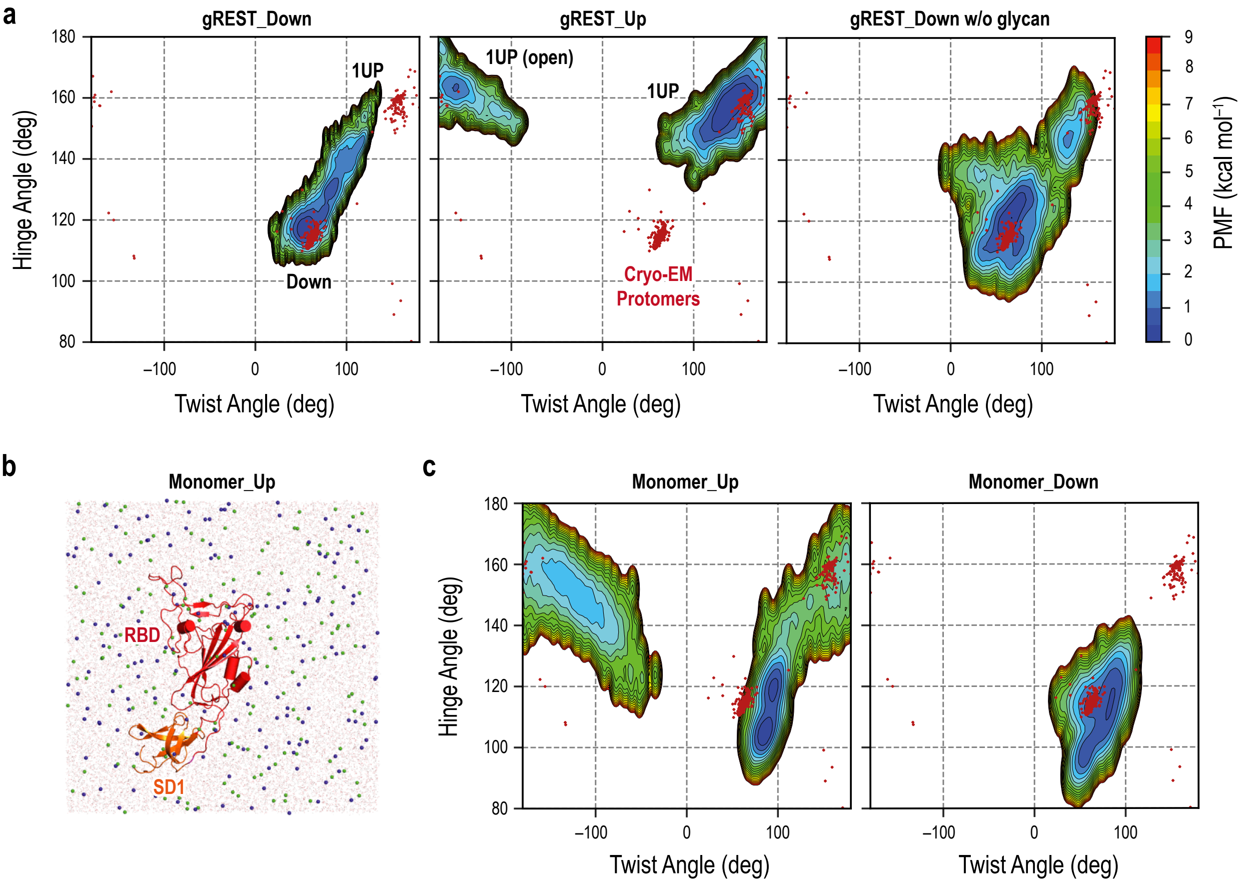


**Fig. S8: Free energy landscape (FEL) along the Hinge/Twist angles in the gREST_SSCR simulations in comparisons with the cMD simulations for RBD/SD1 monomer and cryo-EM structures. a)** FEL of $RBD_A$ in all three gREST_SSCR simulations. Note that these FELs are identical with those in Fig. S6d (left), S7c (left), and S7b (left). 373 Cryo-EM protomers are shown as red dots. This reflects the flexibility of RBD not only in simulations but also in Cryo-EM structures. **b)** Snapshot at 100 ns in the cMD simulation of RBD/SD1 monomer, where the system is solvated with 0.15 M NaCl solution. **c)** FEL of RBD/SD1 monomer starting from Up (left) and Down (right) conformations, highlighting the flexibility of RBD in the absence of protein environment.

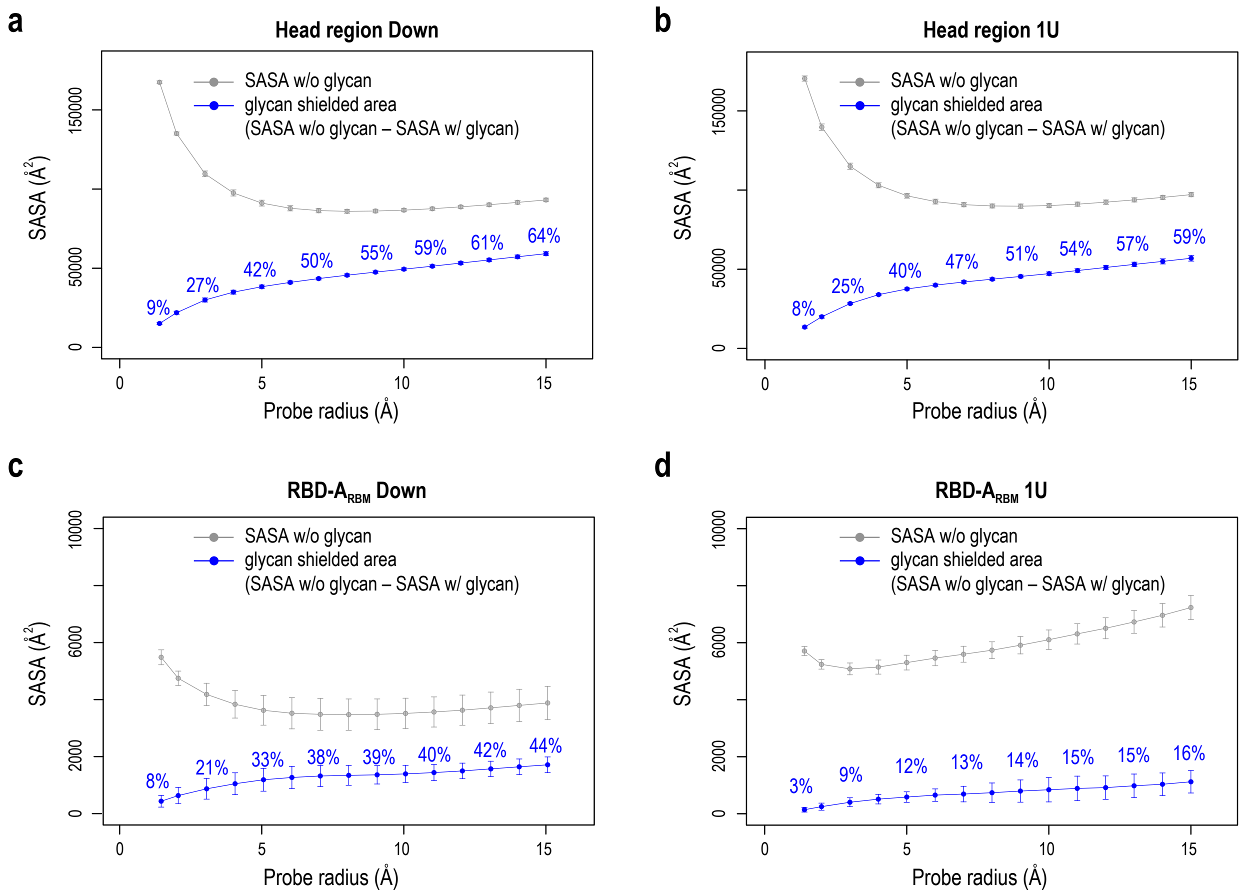


**Figure S9. Glycan-shield of Spike (S) protein. a, b)** Solvent accessible surface area (SASA) of the head region of S protein and the glycan shielded area, calculated for Down and 1U conformations at different probe radii from 1.4Å to 15Å (from a sphere of water to that of antibody scale). The subtraction of the glycan shielded area (blue) from SASA without glycan (gray) gives SASA with glycan (an area between gray and blue curves). The ratio (%) of the glycan shielded area over SASA without glycan is also shown. **c, d)** SASA of the receptor binding motif (RBM, residues 410 to 510) and the glycan shielded area, calculated for Down and 1U conformations. The results are consistent with the previous work by Amaro and co-workers[27].

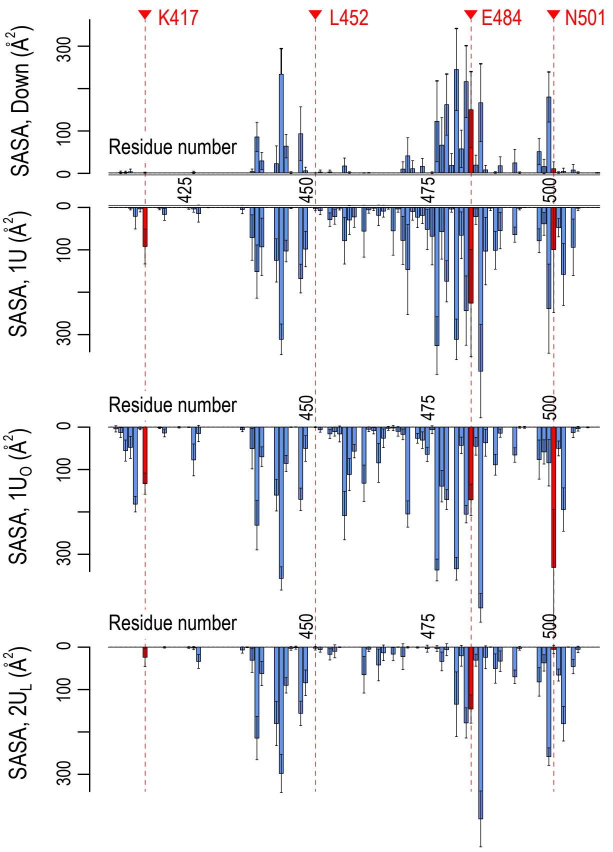


**Figure S10. Accessibility of receptor binding motif (RBM).** Per-residue solvent accessible surface area (SASA) values of the receptor binding motif (RBM, residues 410 to 510) in Down (top) and three Up conformations (1U, 1U$_O$, and 2U$_L$, bottom three). SASA values were calculated using the probe radius of 7.2 Å. Four mutational residues, K417, L452, E484, and N501, are highlighted in red.

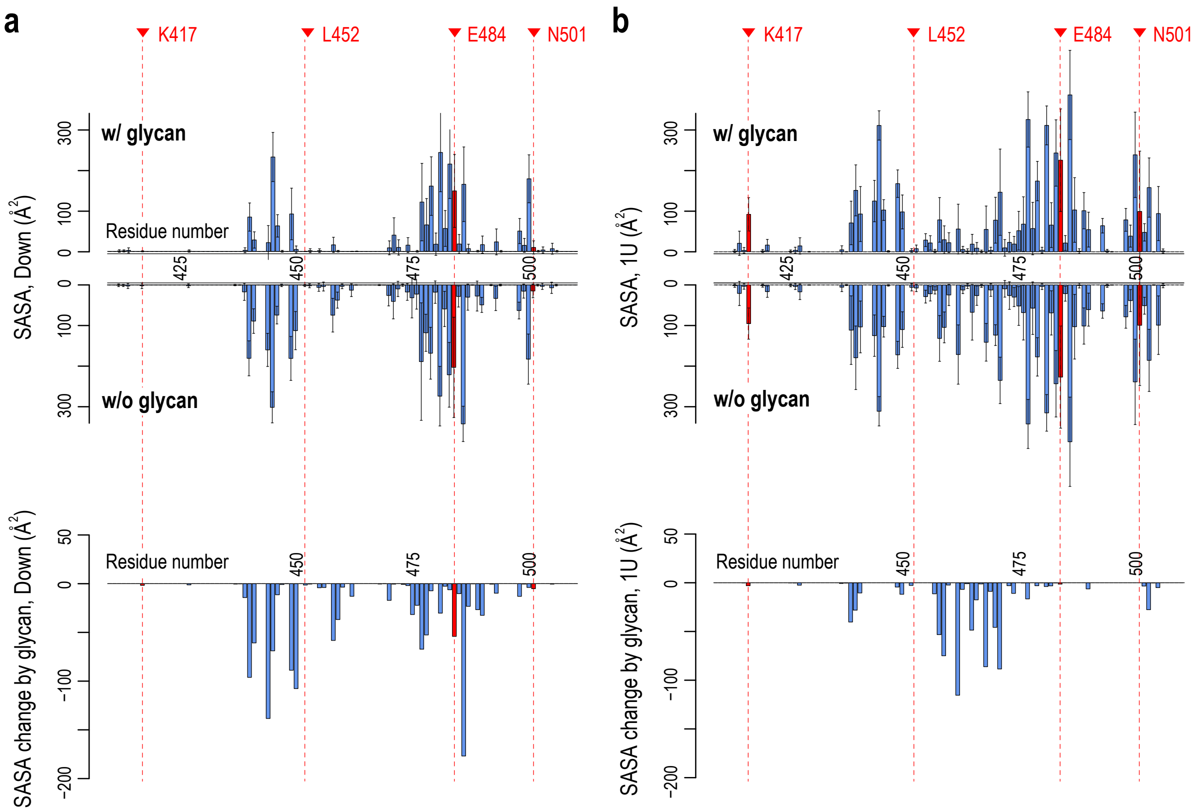


**Figure S11. Glycan effect on the accessibility of receptor binding motif (RBM). a)** Per-residue solvent accessible surface area (SASA) values of the receptor binding motif (RBM, residues 410 to 510) in Down conformation with and without glycan (top) and their changes (SASA w/ glycan – SASA w/o glycan, bottom). b) Per-residue RBM SASA values in 1U conformation with and without glycan and their changes. Four mutational residues, K417, L452, E484, and N501, are highlighted in red.

**a**

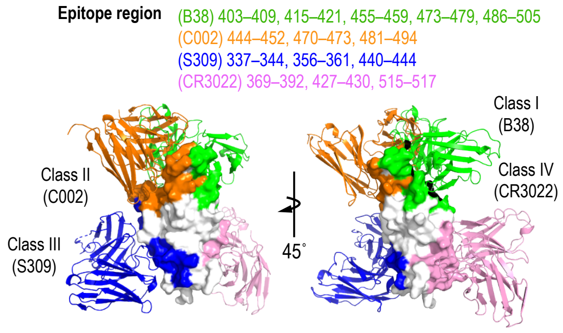


**c**

Crash score in antibody (aligned with "up" RBD) – S-protein pairs

Number of overlaping Cα atoms (< 5Å distance) between the aligned antiboy and S-protein

| | S309 (Class III) PDB: 6WPT | C002 (Class II) PDB: 7K8T | C105 (Class I) PDB: 6XCM |
|---|---|---|---|
| **1U** | 0 | 0 | 2 |
| **$1U_O$** | 11 | 0 | 0 |
| **$2U_L$** | 0 | 0 | 6 |

**b**

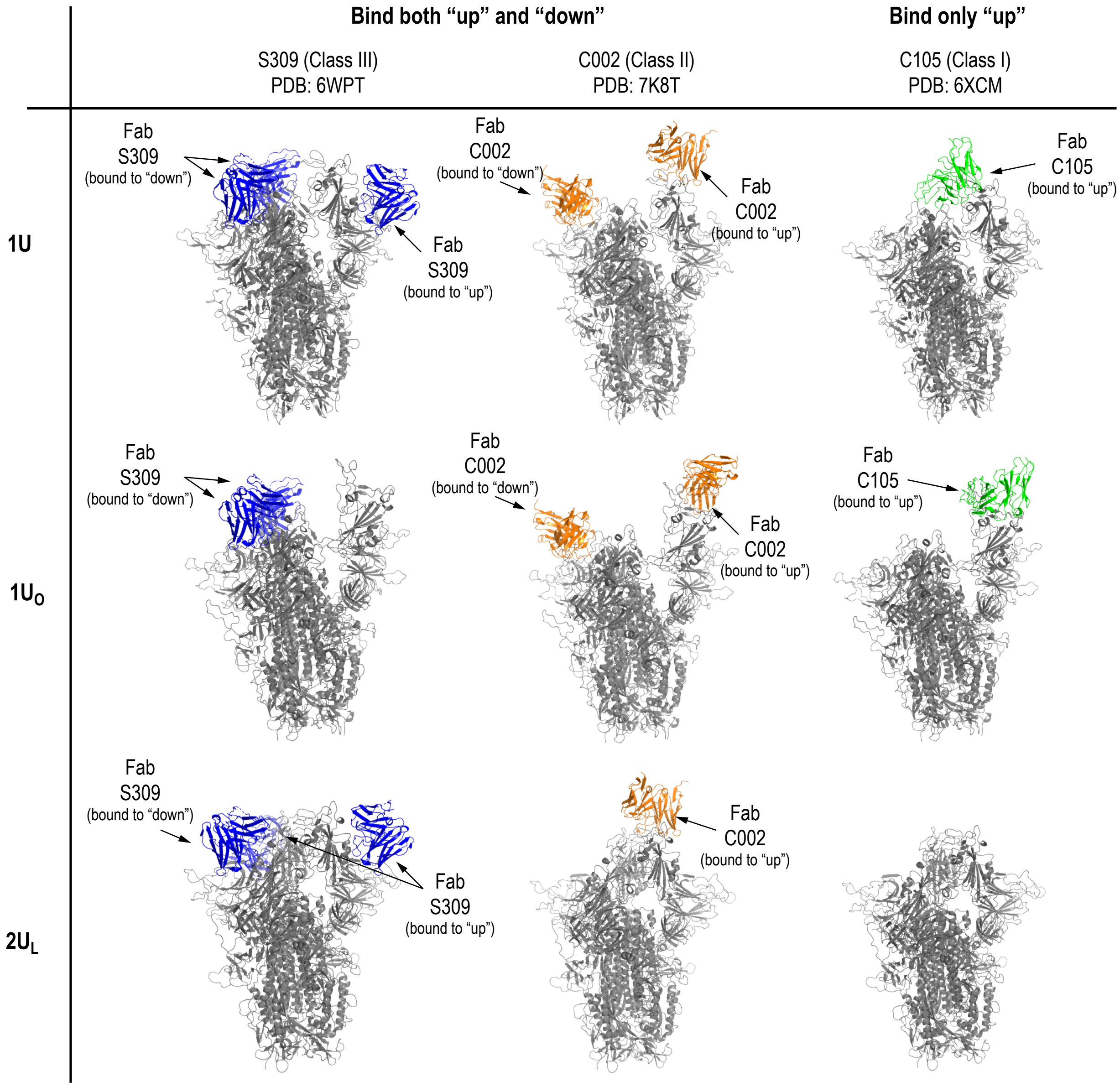


**Figure S12. Putative interaction models with antibodies. a)** The surface representation of RBD epitopes for neutralizing antibodies: green: B38 (Class I), orange: C002 (Class II), blue: S309 (Class III), and pink: CR3022 (Class IV). The structures of S-protein bound antibodies are also shown in cartoon representation. The structures (PDBIDs 6WPT, 7K8T, and 6XCM for S309, C002, and C105, respectively) and epitope residues were taken from the paper by Barnes and co-workers[30]. **b)** The putative interaction with antibodies modeled by aligning the antibody structures from Cryo-EM with each of 1U, $1U_O$, and $2U_L$ conformations. The structures of three antibodies, S309 (Class III), C002 (Class II), and C105 (Class I), were aligned with each conformation. Note that the class II/III antibodies bind both "up" and "down" conformations, while class I binds only "up" conformation. For example, in 1U, S309 and C002 also bind to the "down" RBDs when sterically allowed (In 1U, two "down" RBDs bind S309, while one "down" RBD binds C002 due to steric hindrance). **c)** The crash score between the antibody aligned with "up" RBD and S-protein. We defined the crash score as the number of overlapping Cα atoms (< 5 Å distance) between the aligned antibody and S-protein. To avoid the confusion, the notation of "up" and "down" are used to represent RBD conformation in general.

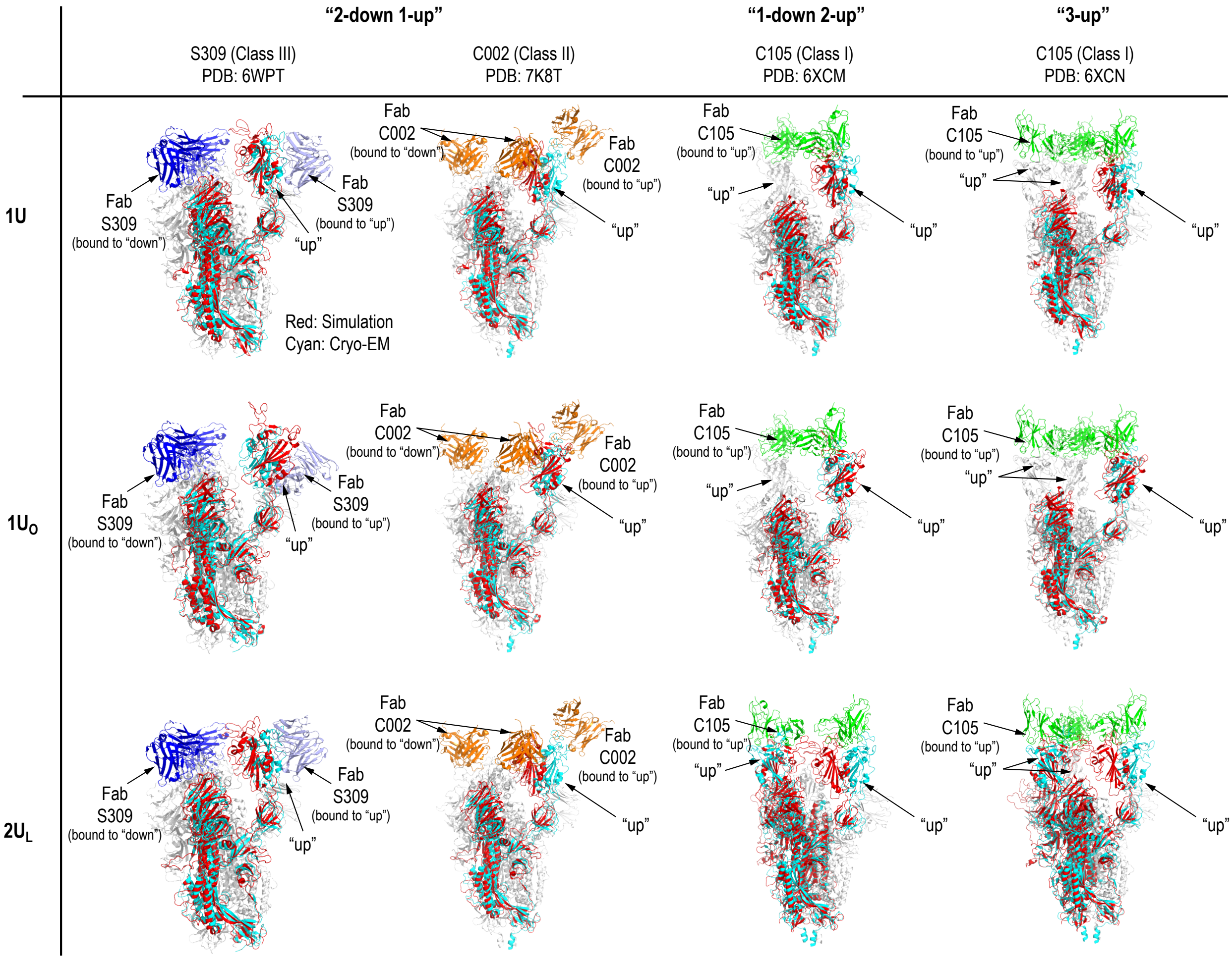


**Figure S13. Comparison of Up structures from MD simulations with Cryo-EM structures.** The structures of 1U, 1U$_O$, and 2U$_L$ from our simulations were aligned with the Cryo-EM structures of S-protein complexed with three types of neutralizing antibodies (nAbs): Class III (S309-S complex, PDBID: 6WPT), Class II (C002-S complex, PDBID: 7K8T), and Class I (C105-S complex, PDBID: 6XCM and 6XCN). The chain A, which involves "up" RBD, of the simulation structure was aligned to the corresponding chain of the Cryo-EM structures using Cα atoms. The aligned structures were colored in red and the corresponding chain of Cry-EM structures were in cyan. Note that Class I nAbs bind only "up" RBD, while Class II/III nAbs bind both "up" and "down" RBDs. In the Cryo-EM structure of S309-S complex, nAb bound to "up" RBD was not resolved, thus the S309 orientation was modeled using the S309-RBD structure resolved for "down" RBD (colored in light blue). To avoid the confusion, the notation of "up" and "down" are used to represent RBD conformation in general.

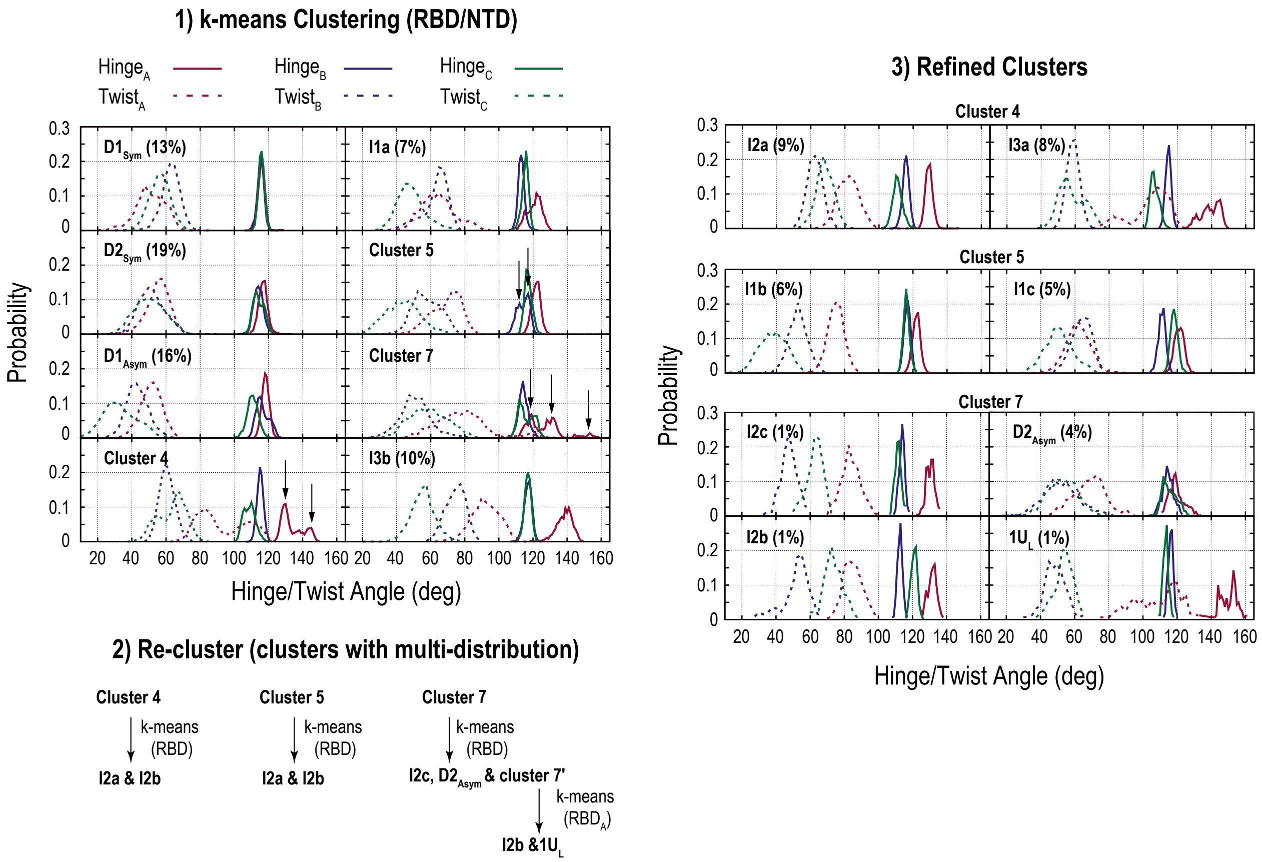


**Figure S14: Clustering for the conformations obtained from the gREST_Down simulation.** Schematic representation of the clustering steps and the resultant distributions of the Hinge and Twist angles are illustrated. 1) First, all conformations at 310 K were classified into 8 clusters using the k-means clustering algorithm, upon fitting S2 and selecting RBD and NTD. Then, the distributions of the Hinge/Twist angles in each cluster were examined. Here, clusters that had multiple peaks are highlighted by black arrows. 2) The clusters with multiple distributions (4, 5 and 7) were further classified into two or three clusters, where only RBD was used as a clustering criterion. 3) The obtained clusters were further examined, and cluster7` was classified into two clusters. Finally, the Hinge/Twist angle distributions in all refined clusters are shown in the right panel.

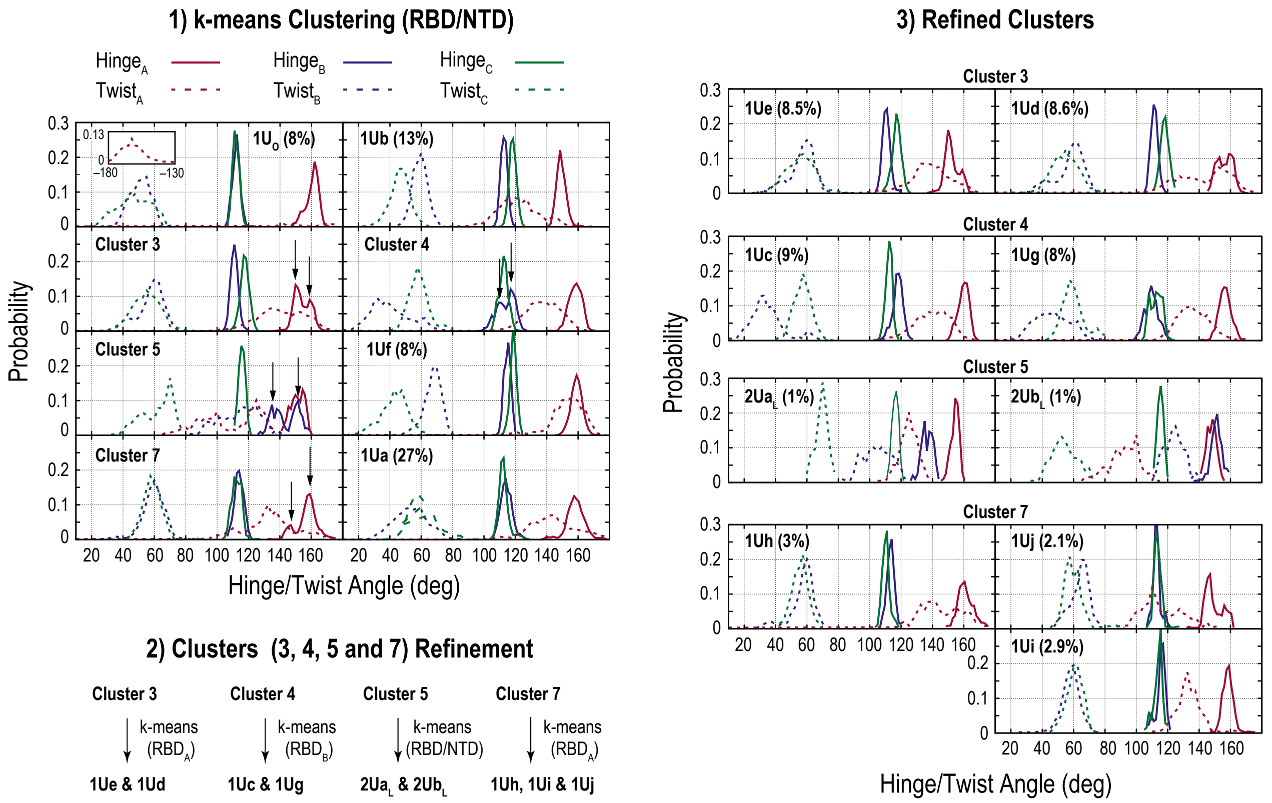


**Figure S15: Clustering for the conformations obtained from the gREST_Up simulation.** The scheme is almost same as in Figure S14.

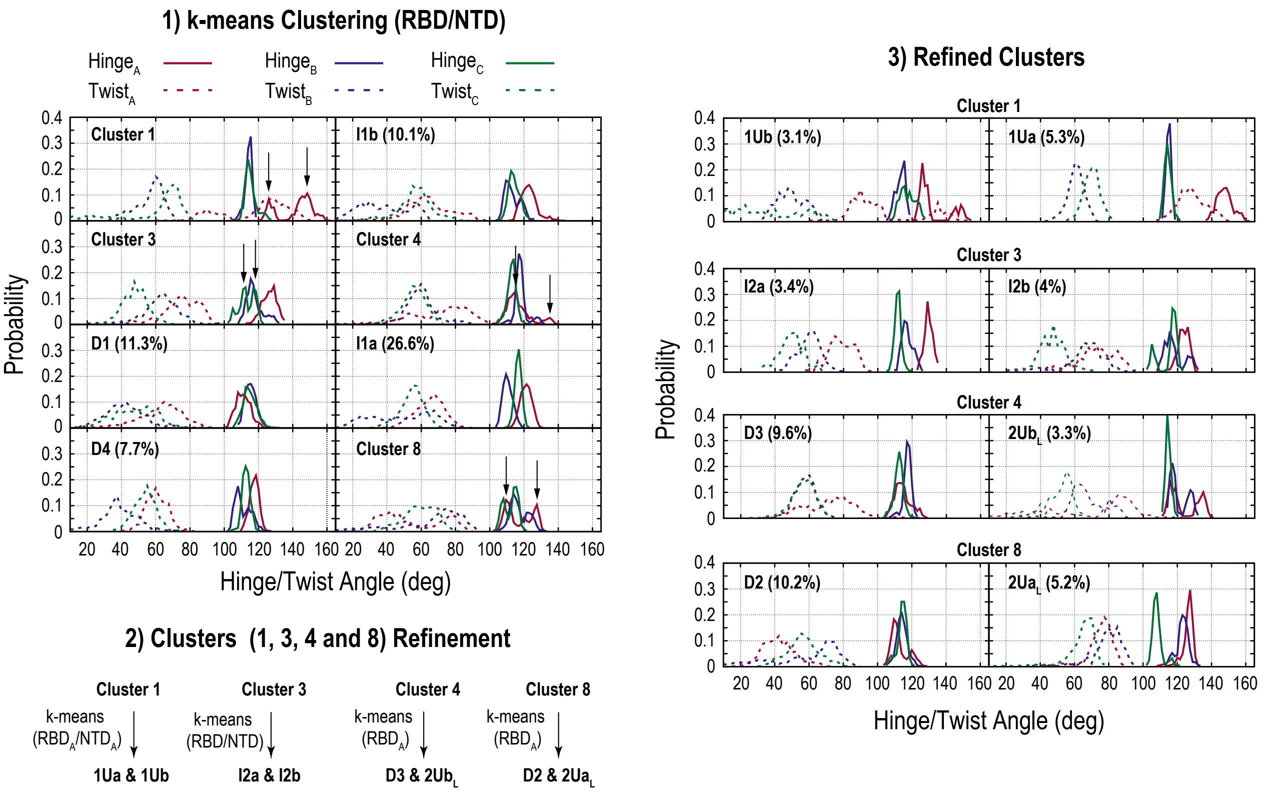


**Figure S16: Clustering for the conformations obtained from the gREST_Down w/o glycan simulation.** The scheme is almost same as in Figure S14.

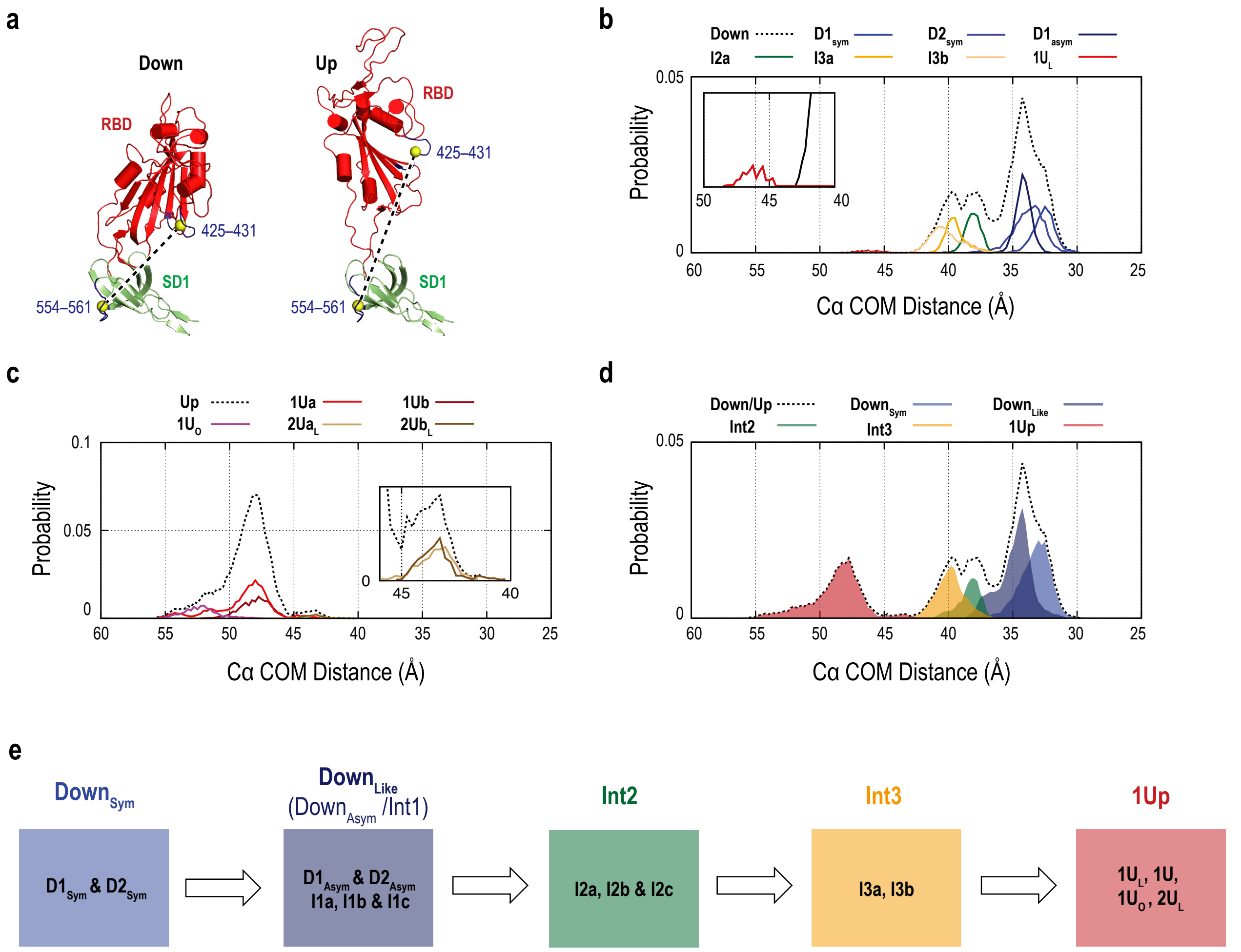


**Figure S17. Simulated smFRET distance using the gREST_SSCR trajectory data. a)** The residues used in the calculation of the smFRET-like distance are illustrated in blue. The distance was estimated based on the center of mass (COM) of the Cα atoms of the residues 425–431 in RBD and 554–561 in SD1 (yellow spheres). **b)** Probability distribution of the smFRET-like distance (black dashed line) in the conformations obtained at 310 K in the gREST_Down simulation. Contributions from main clusters [$Down_{Sym}$ ($D1_{Sym}$ and $D2_{Sym}$), $Down_{Asym}$ ($D1_{asym}$), Intermediates 2 and 3 (I2a, I3a and I3b), and 1Up like ($1U_L$) conformations] are also shown. **c)** Probability distribution of the smFRET-like distance (black dashed line) in the gREST_Up simulation. Main clusters include 1Ua, 1Ub, $IU_O$, 2Ula, and 2Ulb are also shown. **d)** Probability distribution of the smFRET-like distance, where the gREST_Down (b) and gREST_Up (c) were combined with the experimental statistical ration of 77 and 23%. The distribution of macro-clusters that align with the experiment data are also shown. e) Schematic representation of the constituent of the macro-clusters used in (d). For complete description of the formation of macro-clusters from micro-clusters see Table S3.

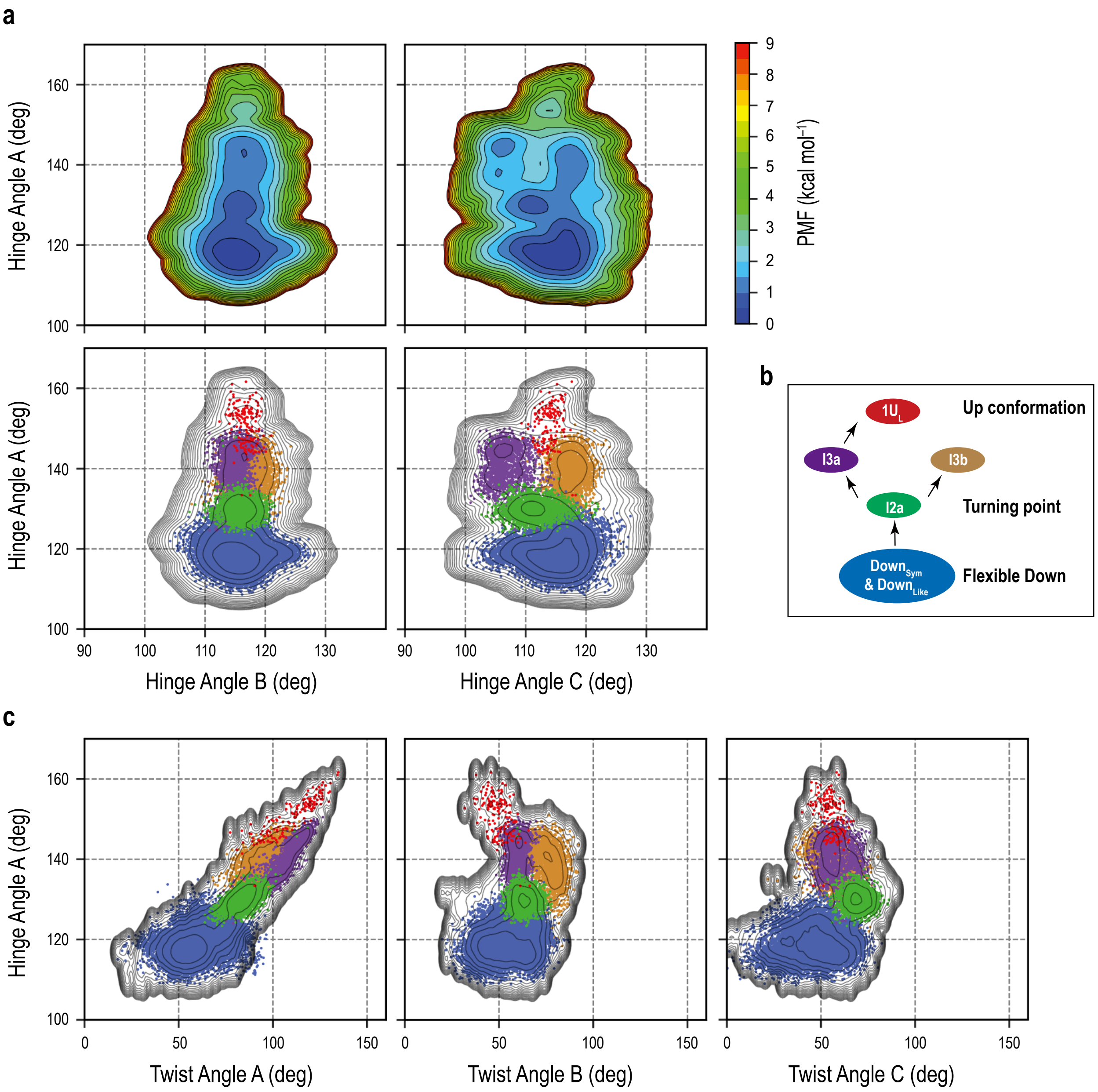


**Figure S18: Transition pathway from Down to 1Up in the gREST_Down simulation. a)** Top: FEL along the $Hinge_A$/ $Hinge_B$ and $Hinge_A$/ $Hinge_C$ angles, and Bottom: projection of the five main clusters [$Down_{Sym}$ & $Down_{Like}$ (blue), I2a (green), I3a (purple), I3b (brown), and $1U_L$ (red)] onto the FEL. The results suggest an independent motion of $RBD_A$ from other RBD Hinge motions during the conformational transition from Down to Up. **b)** Proposed transition pathway from flexible Down to I2a then I3a and finally $1U_L$. **c)** Projection of the five main clusters onto the $Hinge_A$/$Twist_A$, $Hinge_A$/$Twist_B$ and $Hinge_A$/$Twist_C$ maps.

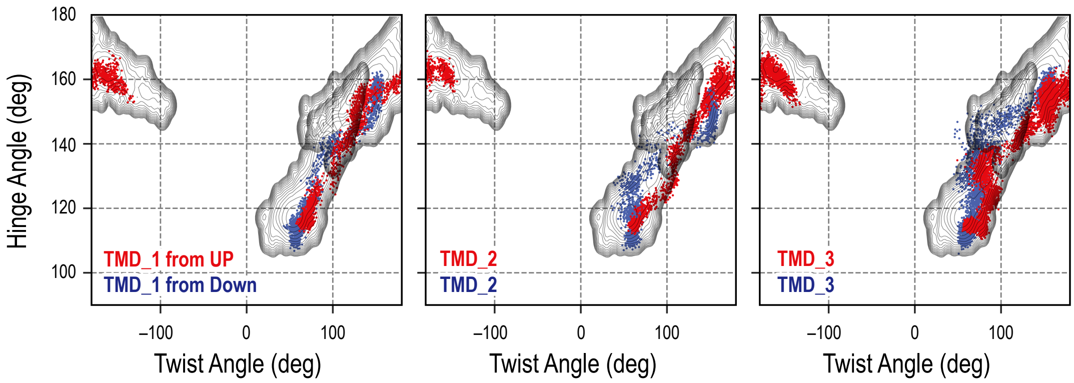


**Figure S19: Comparison between TMD and gREST_SSCR simulations.** Projection of our previous targeted MD (TMD) [Down to Up (blue) or Up to Down (red) simulations[29]] onto the overlapped free energy landscape along the Hinge/Twist angles in the gREST_Down and gREST_Up simulations. In TMD, the simulation time is 50 ns for TMD_1, and 20 ns for TMD_2 and TMD_3, where the different random seeds were used.

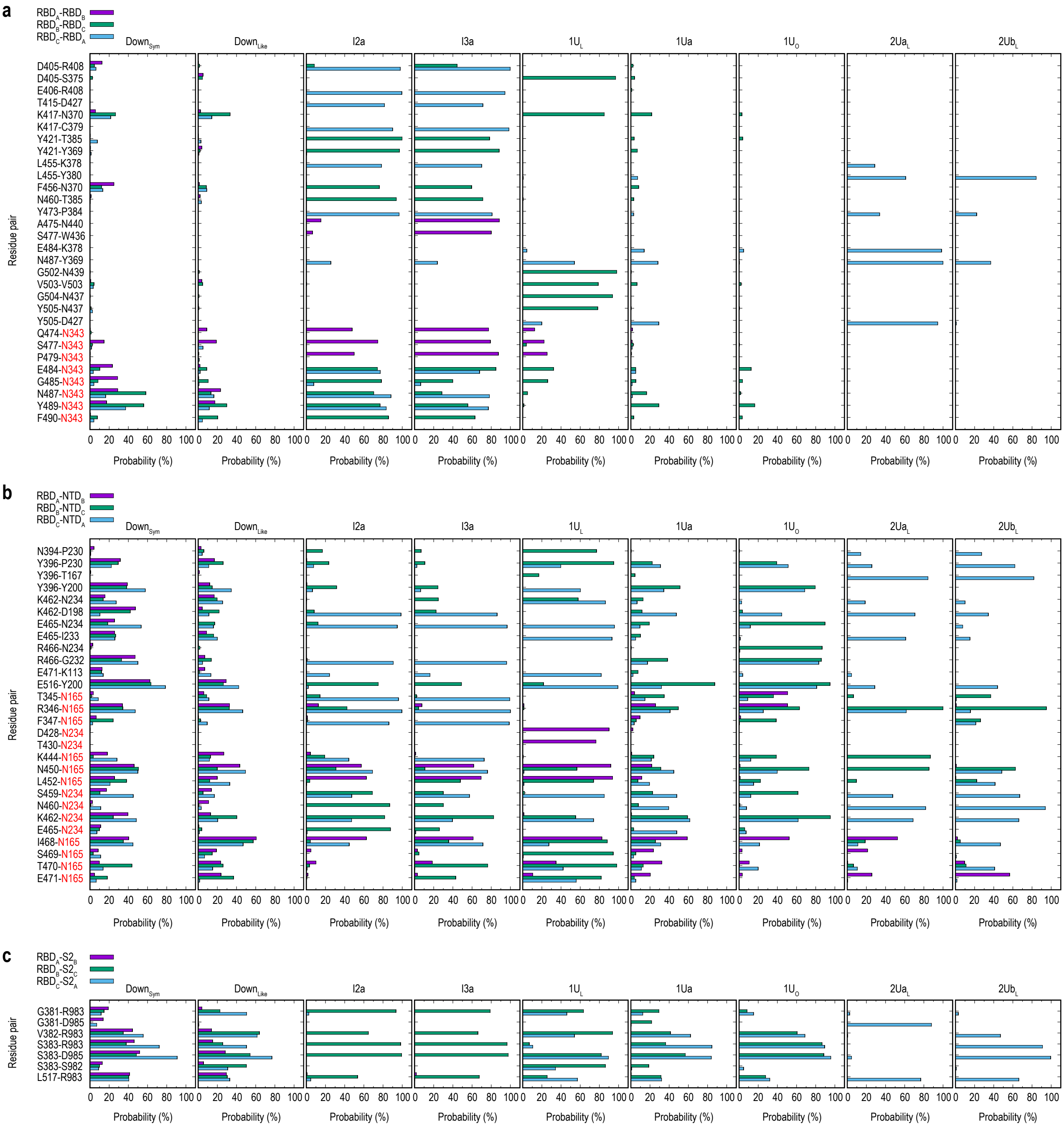


**Figure S20: Contact analysis for the main clusters in gREST_Down and gREST_Up simulations.** Probability of the residue-residue contacts in the RBD/RBD (a), RBD/NTD (b), and RBD/S2 interfaces (c) were analyzed for the main clusters $\mathrm{Down_{Sym}}$ ($\mathrm{D_{Sym}}$), $\mathrm{Down_{Like}}$ ($\mathrm{D_{Asym}}$ and Int1), I2a, I3a, $\mathrm{1U_L}$, top populated 1Up cluster (1Ua), 1Up/open conformation ($\mathrm{1U_O}$), and 2Up like conformation ($\mathrm{2Ua_L}$ and $\mathrm{2Ub_L}$). Contact pairs are selected based on a minimum of 75% probability in any clusters. Pair that involve protein-glycan interactions are highlighted with red color for the glycan part. **a)** Contacts between $\mathrm{RBD_A/RBD_B}$, $\mathrm{RBD_B/RBD_C}$, and $\mathrm{RBD_C/RBD_A}$ are shown in purple, green, and blue respectively. **b)** Contacts between RBD/NTD, where the same colors are used to show $\mathrm{RBD_A/NTD_B}$, $\mathrm{RBD_B/NTD_C}$ and $\mathrm{RBD_C/NTD_A}$ respectively. **c)** Contacts between RBD/S2 showing $\mathrm{RBD_A/S2_B}$, $\mathrm{RBD_B/S2_C}$, and $\mathrm{RBD_C/S2_A}$ in purple, green, and blue respectively.

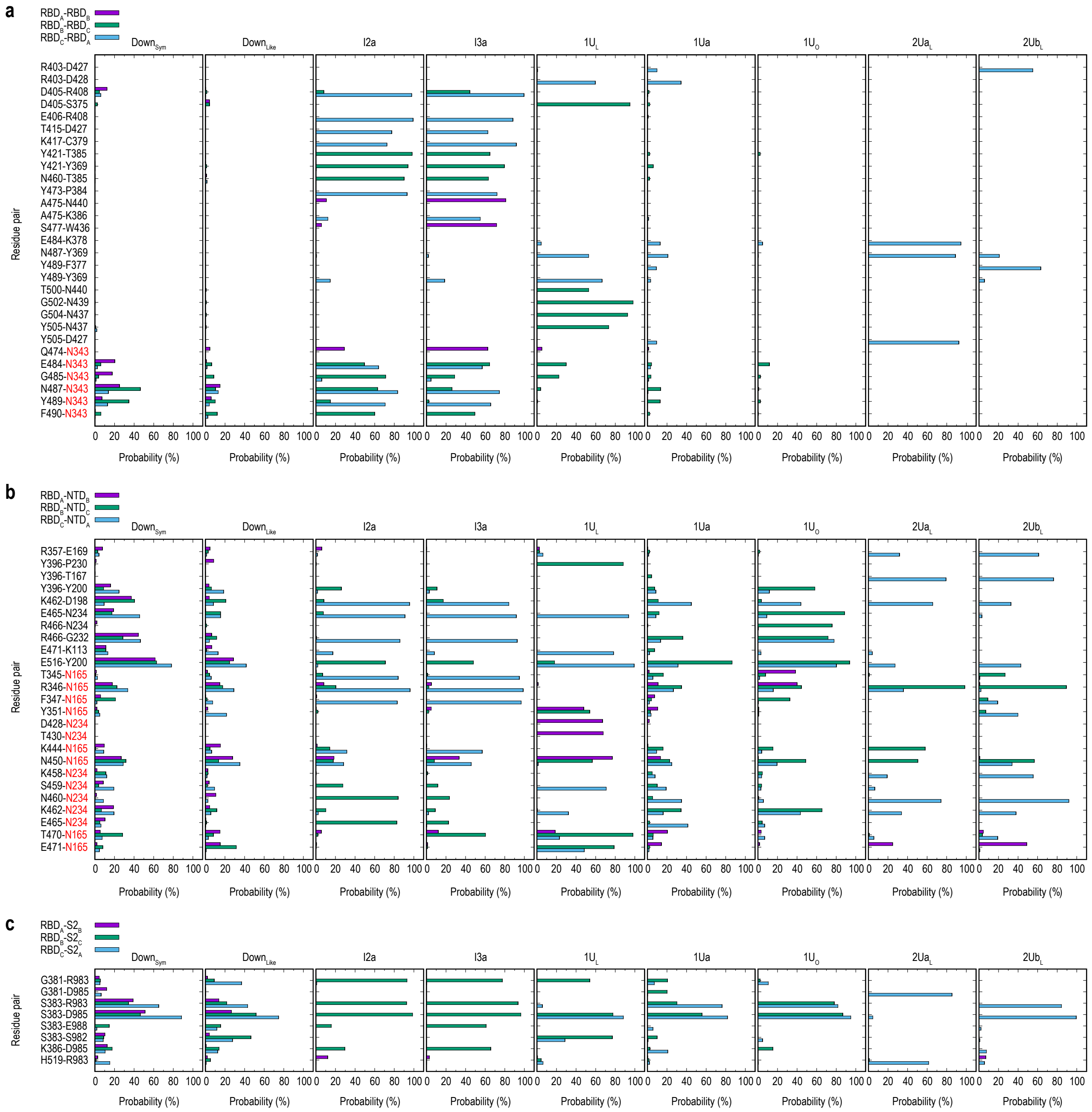


**Figure S21: Hydrogen bond analysis for the main clusters in gREST_Down and gREST_Up simulations.** Probability of the residue-residue hydrogen-bonding in the RBD/RBD (a), RBD/NTD (b), and RBD/S2 interfaces (c) were analyzed. Hydrogen bonding pairs are selected based on a minimum of 50% probability in any clusters. The color definitions are same as in Figure S21.

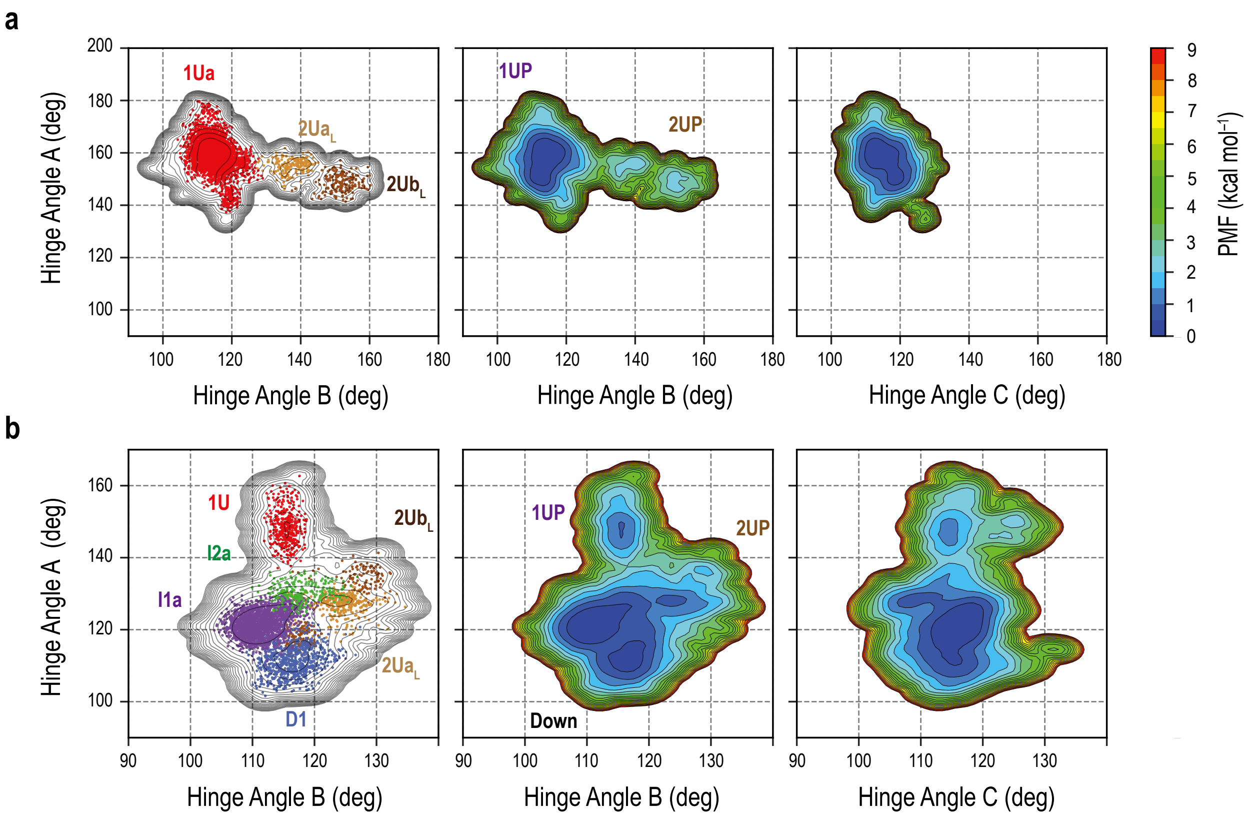


**Figure S22: Free energy landscape in the gREST_Up and gREST_Down w/o glycan simulations. a)** FEL along the $Hinge_A$/$Hinge_B$ (middle) and $Hinge_A$/$Hinge_C$ (right) as well as the projection of the top populated clusters [1Up (1Ua) and 2Up like clusters ($2Ua_L$ and $2Ub_L$)] onto the FEL (left) in the gREST_Up simulation are illustrated. **b)** FEL in the gREST_Down w/o glycan along $Hinge_A$/$Hinge_B$ (middle) and $Hinge_A$/$Hinge_C$ (right). Main clusters along Down to 1Up and Down to 2Up-like conformations are shown in left.

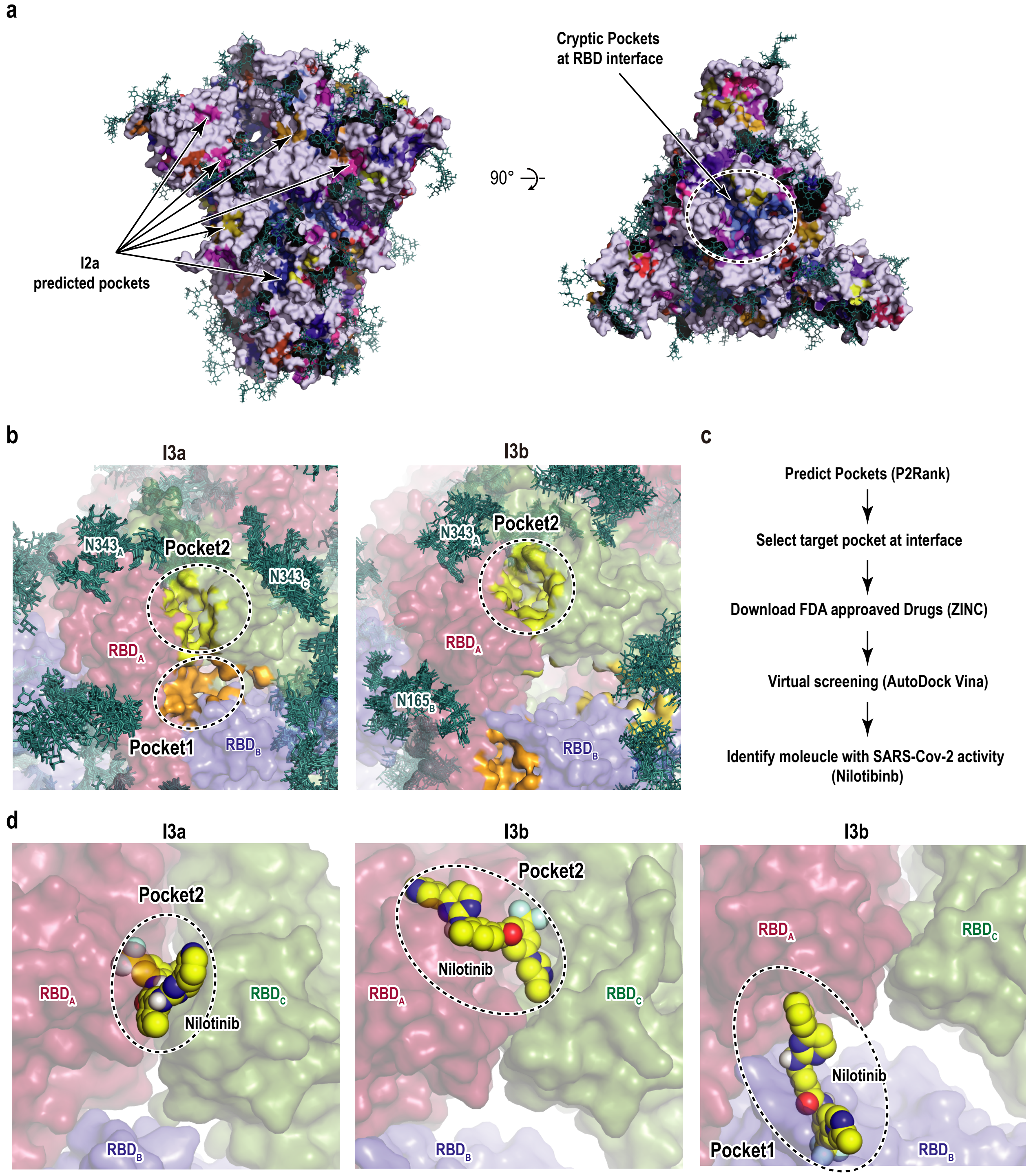


**Fig. S23: Cryptic pockets and ligand binding in RBD. a)** Results of the binding pocket search for I2a predicted by P2Rank software[28]. The spike protein is shown as grey surface, while all other colors represent predicted pockets. Glycans are shown as deep teal sticks, and cryptic pockets at the RBD interface are highlighted by black circle in the top view (right). **b)** Cryptic pockets at the RBD interface in the main intermediate clusters I3a and I3b. $RBD_A$, $RBD_B$, and $RBD_C$ are shown as red, blue, and green surfaces, respectively, and Pocket1 and Pocket2 are shown as yellow and orange surfaces, respectively. Glycans from 10 different conformations that were close to the cluster centers are shown as deep teal sticks. **c)** Scheme for the identification of cryptic pockets and virtual screening used in this study. **d)** Binding of Nilotinib to the predicted cryptic pockets in I3a and I3b. Nilotinib is shown as the sphere model with yellow carbon atoms.

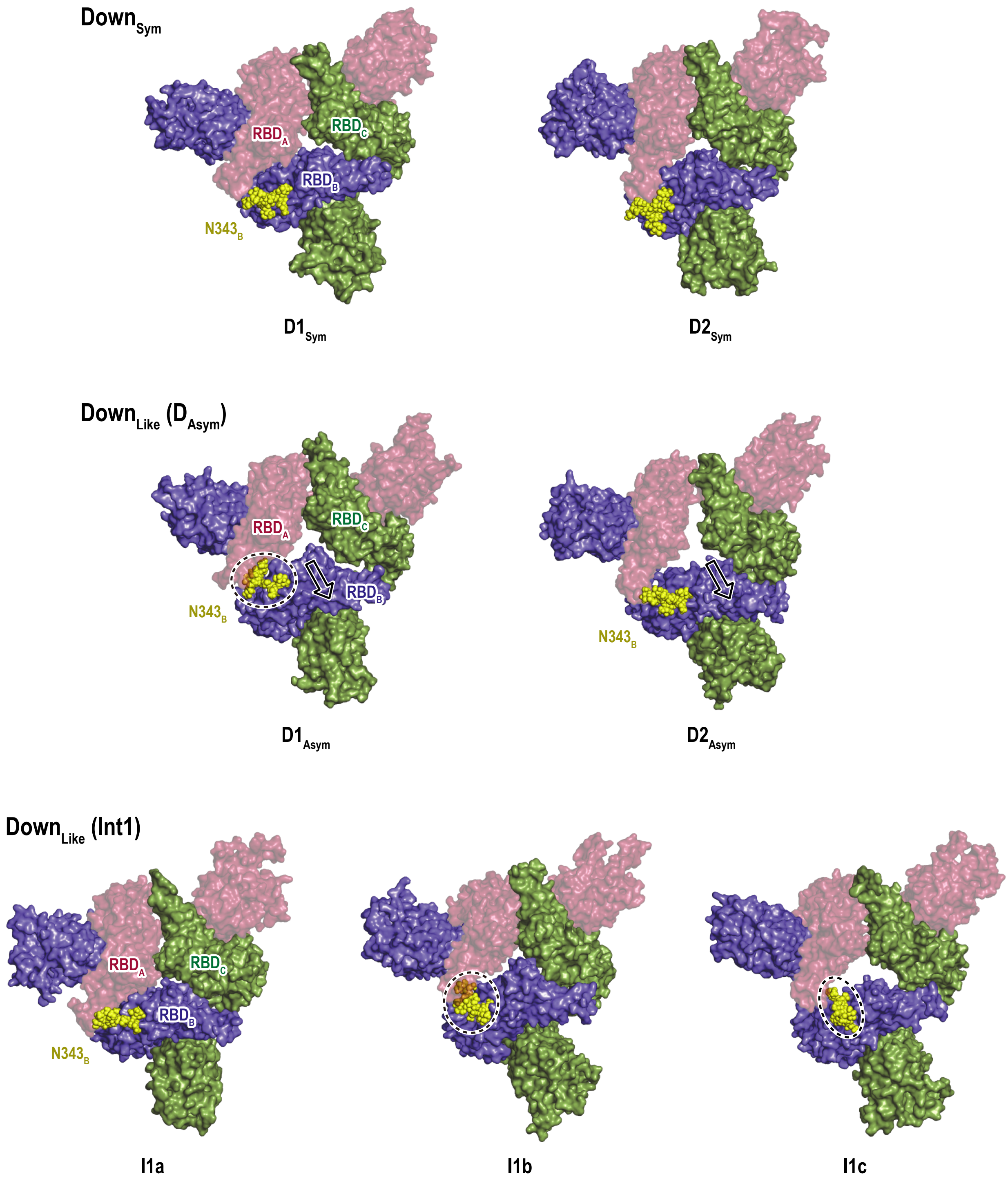


**Figure S24: Relationship between the sideway motion of RBD$_B$ and intrusion of the glycan at N343$_B$.** Cluster centers of Down$_{Sym}$ and Down$_{Like}$ (D$_{Asym}$ and Int1) conformations are shown to highlight the sideway motion of RBD$_B$ that allows the glycan N343 (yellow sphere) to intrude underneath RBD$_A$. Black circle highlight the change in N343$_B$ position.

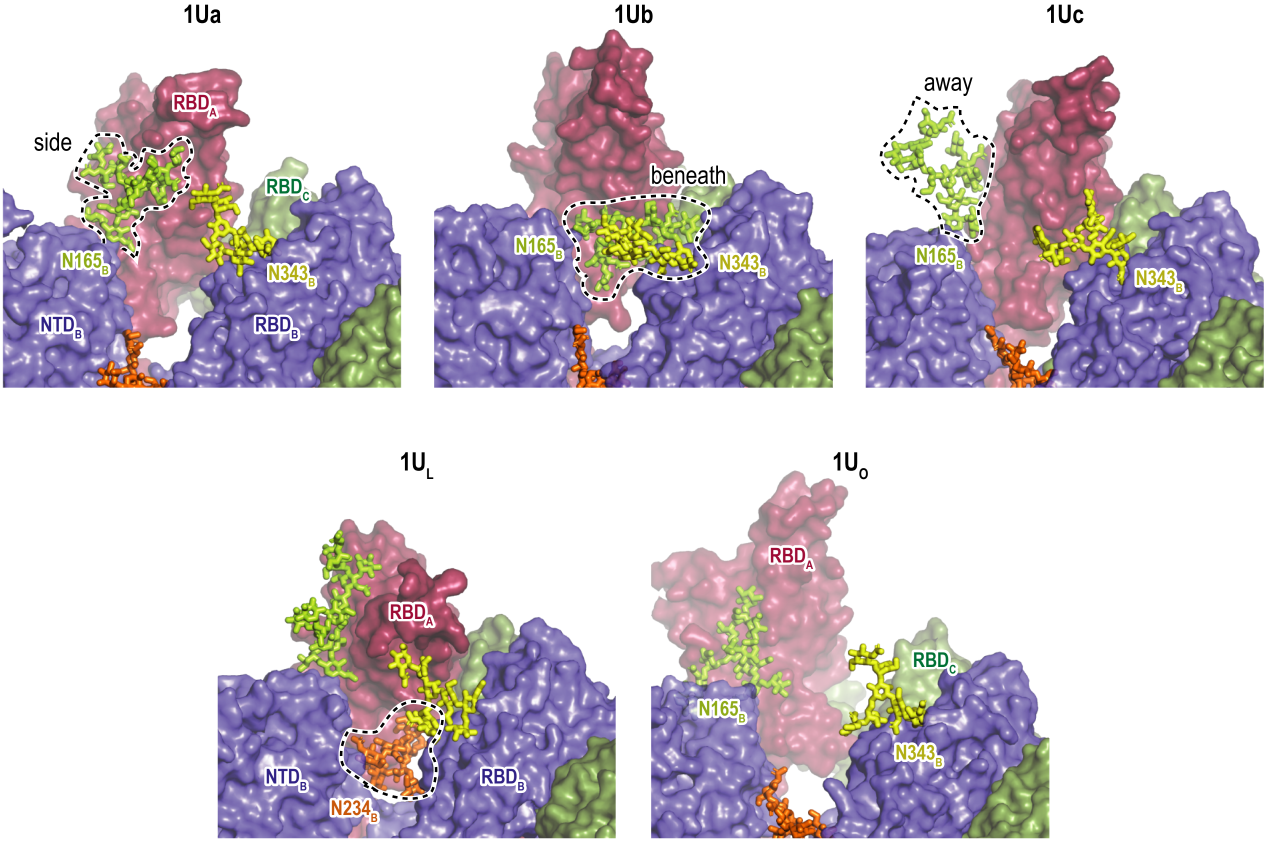


**Fig. S25: Glycan interaction sites in various Up conformations.** Structures of the cluster centers of top populated 1Up (1Ua, 1Ub, and 1Uc) and 1Up/open conformations ($1U_O$) in the gREST_Up simulation are shown. For comparison, the cluster center of 1Up_like ($1U_L$) in the gREST_Down simulation is also shown. Three glycans $N165_B$ (lime), $N234_B$ (orange) and $N343_B$ (yellow) are shown with the stick model, highlighting the diversity of glycan interactions in Up conformations.
